# Split-wrmScarlet and split-sfGFP: Tools for faster, easier fluorescent labeling of endogenous proteins in *Caenorhabditis elegans*

**DOI:** 10.1101/2020.07.02.185249

**Authors:** Jérôme Goudeau, Catherine S. Sharp, Jonathan Paw, Laura Savy, Manuel D. Leonetti, Andrew G. York, Dustin L. Updike, Cynthia Kenyon, Maria Ingaramo

**Affiliations:** Calico Life Sciences LLC, South San Francisco, California 94080, United States; Mount Desert Island Biological Laboratory, Bar Harbor, Maine 04672, United States; Chan Zuckerberg Biohub, San Francisco, California 94158, United States

**Author notes:** Corresponding authors: Calico Life Sciences LLC, 1170 Veterans Blvd., South San Francisco, CA 94080., and.

**Keywords:** *C. elegans*, Cas9, CRISPR, genome engineering, GFP, protein localization, mScarlet, germline

## Abstract

We create and share a new red fluorophore, along with a set of strains, reagents and protocols, to make it faster and easier to label endogenous *C. elegans* proteins with fluorescent tags. CRISPR-mediated fluorescent labeling of *C. elegans* proteins is an invaluable tool, but it is much more difficult to insert fluorophore-size DNA segments than it is to make small gene edits. In principle, high-affinity asymmetrically split fluorescent proteins solve this problem in *C. elegans*: the small fragment can quickly and easily be fused to almost any protein of interest, and can be detected wherever the large fragment is expressed and complemented. However, there is currently only one available strain stably expressing the large fragment of a split fluorescent protein, restricting this solution to a single tissue (the germline) in the highly autofluorescent green channel. No available *C. elegans* lines express unbound large fragments of split red fluorescent proteins, and even state-of-the-art split red fluorescent proteins are dim compared to the canonical split-sfGFP protein. In this study, we engineer a bright, high-affinity new split red fluorophore, split-wrmScarlet. We generate transgenic *C. elegans* lines to allow easy single-color labeling in muscle or germline cells and dual-color labeling in somatic cells. We also describe ‘glonads’, a novel expression strategy for the germline, where traditional expression strategies struggle. We validate these strains by targeting split-wrmScarlet to several genes whose products label distinct organelles, and we provide a protocol for easy, cloning-free CRISPR/Cas9 editing. As the collection of split-FP strains for labeling in different tissues or organelles expands, we will post updates at doi.org/10.5281/zenodo.3993663

## Introduction

Genetically-expressed fluorophores are essential tools for visualizing and quantifying cellular proteins. In *C. elegans*, fluorescent proteins have traditionally been introduced on extrachromosomal arrays (Kimble *et al*. 1982; Mello 1991) or via MosSCI-based integration (Frøkjær-Jensen *et al*. 2008; Frøkjær-Jensen *et al*. 2012). These methods have enabled important discoveries but can also lead to artifacts due to supraphysiological gene-expression levels and lack of endogenous regulatory control. In recent years, the repertoire of *C. elegans* transgenic tools has expanded [see (Nance and Frøkjær-Jensen 2019) for review], particularly due to advances in CRISPR/Cas9 genome-editing technologies (Paix *et al*. 2014; Dickinson and Goldstein 2016). CRISPR/Cas9 allows precise transgene insertion by homology-directed repair (HDR) and can be used to label an endogenous gene at its native locus with a fluorescent protein (Friedland *et al*. 2013; Dokshin *et al*. 2018; Farboud *et al*. 2019; Vicencio *et al*. 2019).

However, relative to CRISPR/Cas9-mediated integration of smaller transgenes, genomic insertion of large DNA fragments like those encoding fluorescent proteins remains a challenge, both because repair with double-stranded templates is less efficient than repair with single-stranded oligodeoxynucleotide donors (ssODN) (Farboud *et al*. 2019), and because of the requirement for cloning to prepare the HDR donor template. Recent methods such as ‘hybrid’ (Dokshin *et al*. 2018) and ‘nested’ (Vicencio *et al*. 2019) CRISPR remove the need for cloning but still require preparation of the DNA template or several rounds of injections and selection of transgenic progeny. As a result, using CRISPR with small ssODN templates is currently faster, easier, cheaper and more efficient than with large templates. In our lab, we routinely make *C. elegans* genome edits with short ssODN with almost guaranteed success. In contrast, in our experience, large edits using double-stranded DNA templates have higher failure rates and are more time-consuming.

Our preferred approach is to combine the utility of full-length fluorescent proteins with the convenience of short genomic edits, by using high-affinity asymmetrically-split fluorescent proteins (Cabantous *et al*. 2004). These fluorophores typically separate a GFP-like protein between the 10^th^ and 11^th^ strands of the beta barrel, splitting it asymmetrically into a large (FP_1-10_) and a small (FP_11_) fragment. The fragments are not individually fluorescent, but upon binding one another, recapitulate the fluorescent properties of an intact fluorophore (Figure 1A). Unlike the low-affinity split fluorescent proteins used in BiFC assays (Hu *et al*. 2002), high-affinity binding between the fragments is critical here. Our preferred approach for tagging a new cellular protein begins with a *C. elegans* strain expressing the large FP_1-10_ fragment in cells of interest, unattached to any cellular protein. This way, only the small FP_11_ fragment (<72 nt) needs to be inserted to tag the target protein, which will only fluoresce in compartments where it can bind the large fragment. These short insertions tend to be faster, easier, and more reliable than inserting a >600 nt full-length fluorescent protein (Paix *et al*. 2015; Prior *et al*. 2017, Dokshin *et al*. 2018; Richardson *et al*. 2018). Therefore, collections of *C. elegans* lines stably expressing the large FP_1-10_ in different tissues are an invaluable resource allowing rapid fluorescent tagging in a cell type of choice. Stable lines with red FP_1-10_ fragments would be especially useful, given *C. elegans*’ substantial autofluorescence in the GFP channel.

**Figure 1.**
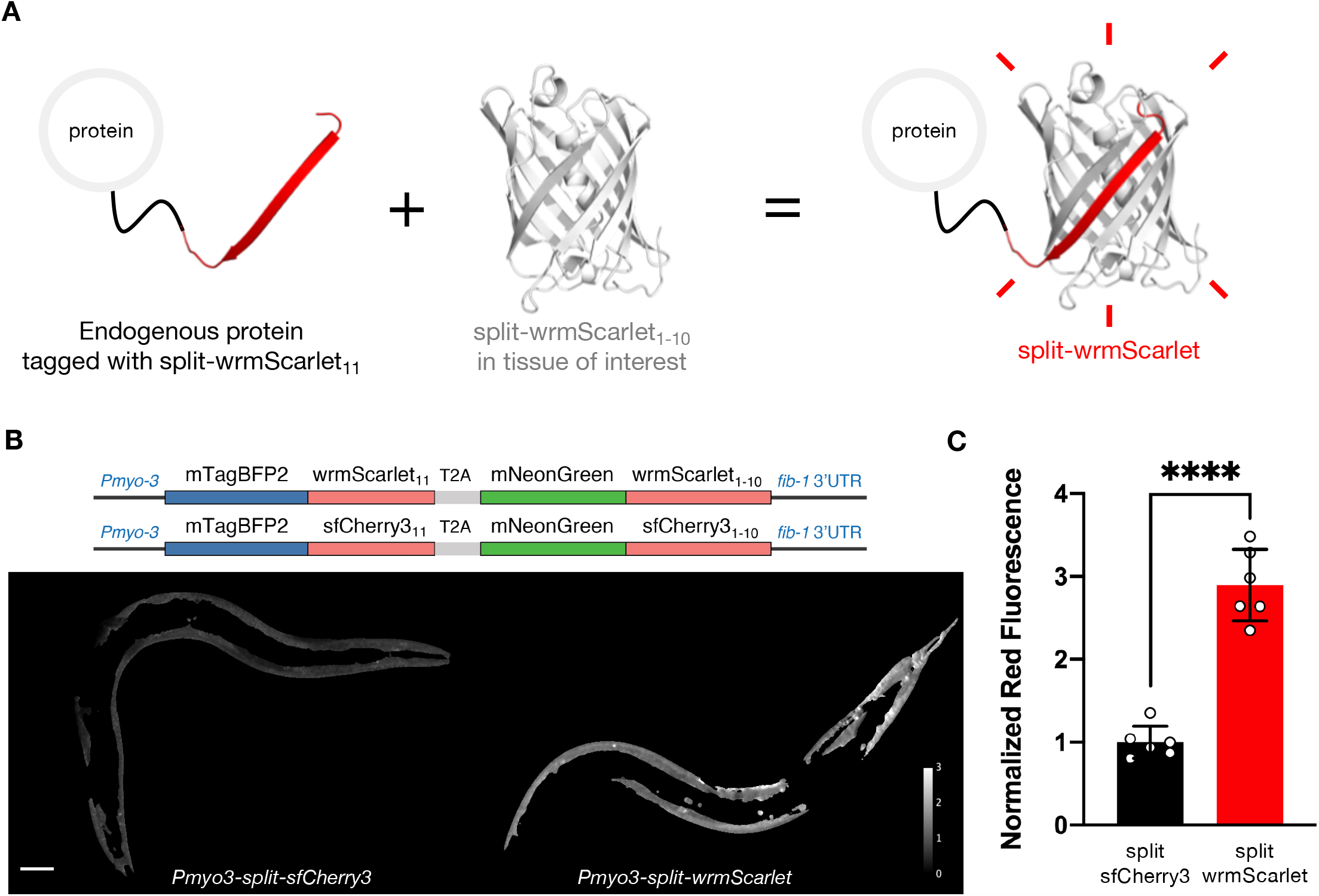
Engineering and evaluating split-wrmScarlet. (A) Principle of endogenous protein labeling with split-wrmScarlet. The protein structure from split-wrmScarlet was generated using Phyre2 and PyMOL. (B) Schematic of the plasmids encoding split-wrmScarlet and split-sfCherry3. Each plasmid consists of the large FP_1-10_ sequence fused to mNeonGreen, and the corresponding small FP_11_ sequence fused to mTagBFP2. The T2A sequence ensures that mTagBFP2::FP_11_ and the corresponding mNeonGreen::FP_1-10_ are separated. The images are representative displays of the ratio of red to green fluorescence intensity from images acquired under identical conditions after background subtraction and masking with the same threshold. Scale bar, 50 µm. (C) Emission intensities from split-sfCherry3 and split-wrmScarlet normalized to mNeonGreen. Mean ± s.d. Circles are individuals (n=6 for each split fluorescent protein). \*\*\*\**P* < 0.0001.

Green and red asymmetrically-split fluorescent proteins have been used to combine cell and protein specificity in *C. elegans* neurons and synapses (Noma *et al*. 2017; He *et al*. 2019; Feng *et al*. 2019); however, these strains used extrachromosomal arrays, not stable lines, which are more time-consuming to maintain and can have variable expression levels. To the best of our knowledge, there is only one available unbound FP_1-10_ stable *C. elegans* line, which expresses sfGFP_1-10_ in the germline (Hefel and Smolikove 2019), and there are no available lines with red FP_1-10_ fragments. Existing red split fluorophores are also much dimmer in *C. elegans* than green ones, despite recent improvements like split-sfCherry3 (Feng *et al*. 2019). In addition, we often struggle to express genome-integrated full-length fluorescent protein fusions in the germline, potentially due to generational silencing.

Here, we describe tools that reduce these obstacles for convenient fluorescent labeling of endogenous *C. elegans* proteins. We engineer split-wrmScarlet, a new split red fluorescent protein based on mScarlet (Bindels *et al*. 2016; El Mouridi *et al*. 2017), which is three times brighter in worms than split-sfCherry3 (https://www.addgene.org/138966). We generate and share *C. elegans* lines carrying single-copy insertions of split-wrmScarlet_1-10_ expressed broadly in somatic cells (https://cgc.umn.edu/strain/CF4582), and specifically in muscle (https://cgc.umn.edu/strain/CF4610). We also describe a novel approach to make *C. elegans* lines with robust germline expression of exogenous proteins that appears to be resistant to generational silencing. We use this approach to make a germline split-wrmScarlet_1-10_ strain (https://cgc.umn.edu/strain/DUP237).

We provide a protocol for an easy, cloning-free method to label endogenous genes with FP_11_s using CRISPR/Cas9, commercially available synthetic single-stranded oligodeoxynucleotide (ssODN) donors, and microinjection (dx.doi.org/10.17504/protocols.io.bamkic4w). We validate this protocol by targeting split-wrmScarlet_11_ to six different genes whose products have distinct cellular locations. We show that labeling with tandem split-wrmScarlet_11_-repeats increases fluorescence *in vivo*, and we provide the plasmid necessary to generate the dsDNA template (https://www.addgene.org/158533). We also generate a strain expressing an integrated copy of sfGFP_1-10_ (Pédelacq *et al*. 2005) broadly in somatic cells (https://cgc.umn.edu/strain/CF4587), and a strain expressing sGFP2_1-10_ (Köker *et al*. 2018) specifically in the germline (https://cgc.umn.edu/strain/DUP223). Finally, we generate a dual-color strain expressing both sfGFP_1-10_ and split-wrmScarlet_1-10_ in somatic cells (https://cgc.umn.edu/strain/CF4588), for two-color applications such as colocalization studies or organelle interaction. We hope that these resources will facilitate the study of *C. elegans* biology. As the collection of split-FP strains and related resources for labeling different tissues, organelles and proteins expands, we will post updates at doi.org/10.5281/zenodo.3993663

## MATERIALS AND METHODS

### Mutagenesis and screening

For the initial screenings in *E. coli*, we introduced a 32 amino-acid spacer between the tenth and eleventh ±-strands of full-length mScarlet in a pRSET vector (Feng *et al*. 2017). This starting construct was non-fluorescent, but we restored low fluorescence levels by introducing the superfolder mutation G220A. Semi-random mutagenesis was carried out using rolling-circle amplification with NNK primers at positions I8, K10, F15, G32, Q43, A45, K46, L47, G52, G53, D60, S63, P64, Q65, F66, S70, R71, T74, K75, D79, Y84, W94, R96, T107, V108, Q110, E115, L125, R126, T128, K139, K140, W144, E145, S147, T148, E149, R150, I162, K163, M164, L175, F178, K179, K183, K185, K186, N195, R198, I202, T203, S204, D208, Y209, T210, V211, V212, E213, Q214, Y215, E216, R217, S218, E219, A220, H222, S223, T224, G225, G226, M227, D228, and E229 with Phusion polymerase (NEB) in GC buffer, followed by pooling of the PCR products, *DpnI* digestion and transformation into BL21(DE3) *E. coli*. These positions covered areas deemed important for brightness or stability, and the interface between FP_11_ and FP_1-10_. Primers were resynthesized if a mutation interfered with neighboring mutagenic primer binding. The brightest three to five colonies were identified using a Leica M165 FC fluorescent stereomicroscope, and their plasmid DNA subjected to a new mutagenesis round. After five rounds, we separated the two fragments of a version of split-wrmScarlet (which had fluorescence comparable to mScarlet) into two *S. cerevisiae* plasmids to test for complementation. Since we did not detect fluorescence, we continued selection using two plasmids in yeast. For screening on two plasmids, a pRSET vector expressing split-wrmScarlet_1-10_ and a pD881-MR vector (ATUM) expressing mTagBFP-split-wrmScarlet_11_ (without the MDELYK tail from the C-terminus) were used to perform the semi-random mutagenesis. The libraries were co-electroporated into *E. coli* and expression was induced with 1% rhamnose and 1mM IPTG. The library was enriched for fluorescent clones using FACS, and then subcloned to make pRS-GPD-split-wrmScarlet_1-10_ and p416-TEF-membrane-mTagBFP-split-wrmScarlet_11_.

The yeast plasmids were co-transformed into a URA^-^, HIS^-^, LEU^-^, MET^-^ *S. cerevisiae* strain and selected for in SC media without uracil and histidine, and FACS was used again for enrichment of clones with the highest red to blue ratio. After three rounds of semi-random mutagenesis with the two-plasmid strategy, a final round of random mutagenesis was performed using the GeneMorph II kit (Agilent). Yeast plasmids are available through Addgene (https://www.addgene.org/158585/, https://www.addgene.org/158584/), and *E. coli* plasmid sequences are present in Supplementary Material, Table S5.

### *C. elegans* strains and maintenance

Animals were cultured under standard growth conditions with *E. coli* OP50 at 20°C (Brenner 1974). Strains generated in this work are listed in the Supplementary Material, Table S3.

### Nucleic acid reagents

Synthetic nucleic acids were purchased from Integrated DNA Technologies (IDT), GenScript or Genewiz. For knock-in of a single split-wrmScarlet_11_ or sfGFP_11_ sequence, 200-mer HDR templates were ordered in ssODN form (synthetic single-stranded oligodeoxynucleotide donors) from IDT. For knock-in of split-wrmScarlet_11_ repeats, HDR templates were ordered in dsDNA form (plasmids) from GenScript or Genewiz. For plasmids injected as extrachromosomal arrays, sequences were synthesized and cloned into the pUC57 vector (Genewiz). The complete set of crRNAs and DNA sequences and plasmids used for the experiments described here can be found in Supplementary Material, Tables S1, S4 and S5.

### Strain generation: CRISPR/Cas9-triggered homologous recombination

CRISPR insertions were performed using published protocols (Paix *et al*. 2015, 2016). Ribonucleoprotein complexes (protein Cas9, tracrRNA, crRNA) and DNA templates were microinjected into the gonad of young adults using standard methods (Evans 2006). Injected worms were singled and placed at 25°C overnight. All crRNA and DNA template sequences used to generate the strains described in this work are listed in the Supplementary Material, Table S4. Split-wrmScarlet_11_ and sfGFP_11_ integrants were identified by screening for fluorescence in the F1 or F2 progeny of injected worms. The co-CRISPR *dpy-10*(*cn64*) mutation was used as a marker when generating non-fluorescent strains. The CF4582 strain (*muIs252*[*Peft-3::split-wrmScarlet*_*1-10*_*::unc-54 3’UTR, Cbr-unc-119*(*+*)] *II; unc-119*(*ed3*) *III*) was generated by replacing the *tir-1::mRuby* sequence from the strain CA1200 (*ieSi57 II; unc-119*(*ed3*) *III* (Zhang *et al*. 2015) with the *split-wrmScarlet*_*1-10*_ sequence. The CF4587 strain (*muIs253*[(*Peft-3::sfGFP*_*1-10*_*::unc-54 3’UTR, Cbr-unc-119*(*+*)] *II; unc-119*(*ed3*) *III*) was generated by replacing the *let-858* promoter from the strain COP1795 (*knuSi785* [*pNU1687*(*Plet-858::sfGFP*_*1-10*_*::unc-54 3 R, unc-119*(*+*)] *II; unc-119*(*ed3*) *III*) with the *eft-3* (also known as *eef-1A*.*1*) promoter. Both CF4582 and CF4587 strains were generated using long, partially single-stranded DNA donors (Dokshin *et al*. 2018). The CF4610 strain (*muIs257*[*Pmyo-3::split-wrmScarlet*_*1-10*_*::unc-54 3’UTR*] *I*) was generated by inserting the split-wrmScarlet_1-10_ sequence in the WBM1126 strain following the SKI LODGE protocol (Silva-García *et al*. 2019). The strains PHX731 (*vha-13*(*syb731*[*wrmScarlet::vha-13*]) *V*) and PHX1049 (*vha-13*(*syb1049*[*gfp::vha-13*]) *V*) were generated by SunyBiotech’s CRISPR services. Strains generated were genotyped by Sanger sequencing of purified PCR products (Genewiz).

### Strain generation: Mos1-mediated single-copy insertion

The COP1795 strain was generated by NemaMetrix’s MosSCI services. The PHX1797 strain was generated by SunyBiotech’s MosSCI services, using a codon-optimized sequence of split-wrmScarlet_1-10_ with three introns, and engineered to avoid piRNA recognition transgene silencing (Wu *et al*. 2018; Zhang *et al*. 2018; Supplementary Material, Table S1).

### Strain generation: genetic crosses

The following *C. elegans* strains were created by standard genetic crosses: CF4588 (*muIs253*[*Peft-3::sfGFP*_*1-10*_*::unc-54 3’UTR, Cbr-unc-119*(*+*)], *muIs252*[*Peft-3::split-wrmScarlet*_*1-10*_*::unc-54 3’UTR, Cbr-unc-119*(*+*)] *II; unc-119*(*ed3*) *III*) and CF4602 (*muIs253*[*Peft-3::sfGFP*_*1-10*_*::unc-54 3’UTR, Cbr-unc-119*(*+*)], *muIs252*[*Peft-3::split-wrmScarlet*_*1-10*_*::unc-54 3’UTR, Cbr-unc-119*(*+*)] *II; unc-119*(*ed3*) *III; fib-1*(*muIs254*[*split-wrmScarlet*_*11*_*::fib-1*]), *his-3*(*muIs255*[*his-3::sfGFP*_*11*_]) *V*). Non-fluorescent parental lines CF4582, CF4587 and CF4610 generated using *dpy-10*(*cn64*) co-CRISPR were backcrossed at least once.

### Strain generation: plasmid microinjection

*Peft-3::3NLS::mTagBFP2::split-wrmScarlet*_*11*_*::T2A::mNeonGreen::split-wrmScarlet*_*1-10*_*::fib-13 R, Peft-3::3NLS::mTagBFP2::sfCherry3*_*11*_*::T2A::mNeonGreen::sfCherry3*_*1-10*_*::fib-1 3 R, Pmyo-3::mTagBFP2::split-wrmScarlet*_*11*_*::T2A::mNeonGreen::split-wrmScarlet*_*1-10*_*::fib-1 3 R*, or *Pmyo-3::mTagBFP2::sfCherry3*_*11*_*::T2A::mNeonGreen::sfCherry3*_*1-10*_*::fib-1 3 R* constructs were microinjected at (20 ng/µL) using a standard microinjection procedure (Mello 1991). Germline gene expression was achieved using a microinjection-based protocol with diluted transgenic DNA (Kelly *et al*. 1997), *Psun-1::mNeonGreen::linker::split-wrmScarlet*_*11*_*::tbb-2 3 R* construct (5 ng/µL) was co-injected with PvuII-digested genomic DNA fragments from *E. coli* (100 ng/µL). Plasmid sequences are listed in Supplementary Material, Table S5.

### Germline strain generation: *glh-1::T2A::split-wrmScarlet*_*1-10*_ *and glh-1::T2A::sGFP2*_*1-10*_

Using CRISPR/Cas9, the C-terminus of *glh-1* was tagged with either T2A::split-wrmScarlet_1-11_ or T2A::sGFP2_1-11,_ a split superfolder GFP variant optimized for brightness and photostability (Köker *et al*. 2018). Fluorescence originating from these full-length fusions was present throughout the cytoplasm and nuclei of adult germ cells and gametes, with the maternally deposited signal persisting through the early stages of embryogenesis and larval development (Figure S8A and S8B, top panels). After verifying fluorescence, we used a precise CRISPR/Cas9 deletion of either split-wrmScarlet_11_ or sGFP2_11_ to convert these FP_1-11_ strains into FP_1-10_ strains, DUP237 *glh-1*(*sam140*[*glh-1::T2A::split-wrmScarlet*_*1-10*_]) *I* and DUP223 *glh-1*(*sam129*[*glh-1::T2A::sGFP2*_*1-10*_]) *I* and corroborated the absence of fluorescence (Figure S8A and S8B, middle panels). The crRNAs, ssDNAs and dsDNA template sequences are described in Supplementary Material, Tables S1-S4.

### Microscopy

Confocal fluorescence imaging was performed using NIS Elements imaging software on a Nikon confocal spinning disk system equipped with an Andor EMCCD camera, a CSU-X1 confocal scanner (Yokogawa), 405, 488, and 561 nm solid-state lasers, and 455/50, 525/26 and 605/70 nm emission filters. Transgenic animals expressing sfGFP_11_ or split-wrmScarlet_11_ were screened using a Leica M165 FC fluorescent stereomicroscope equipped with a Sola SE-V with GFP and mCherry filters.

### Image analysis

Images were analyzed using Fiji. Image manipulations consisted of maximum intensity projections along the axial dimension, rolling ball radius background subtraction, smoothing, and LUT minimum and maximum adjustments. Masks were created by thresholding and setting the pixels under the threshold cutoff to NaN. Plotting of values per pixel was carried out in python 3, using numpy and matplotlib. When performing normalizations for split-sfCherry3 vs split-wrmScarlet, the red channel was divided by the green channel (mNeonGreen::FP_1-10_) because the localization of both fragments is expected to be the same (cytosolic). For normalization of signals where mTagBFP::FP_11_ is targeted to the membrane, the blue channel was used instead of the green channel.

### Mounting worms for microscopy

Pads made of 3% agarose (GeneMate) were dried briefly on Kimwipes (Kimtech) and transferred to microscope slides. Around 10 µL of 2 mM levamisole (Sigma) was pipetted onto the center of the agarose pad. Animals were transferred to the levamisole drop, and a coverslip was placed on top before imaging.

### Brood size analysis

Eight single synchronized adults grown at 20°C were transferred to fresh plates every 24 hours until cessation of reproduction, and the number of viable progeny produced by each worm was scored.

### Developmental toxicity assay

Ten N2E wild-type animals were microinjected with either *Peft-3::3NLS::mTagBFP2::split-wrmScarlet*_*11*_*::T2A::mNeonGreen::split-wrmScarlet*_*1-10*_*::fib-1 3 R* or *Peft-3::3NLS::mTagBFP2::sfCherry3*_*11*_*::T2A::mNeonGreen::sfCherry3*_*1-10*_*::fib-1 3 R* construct at (20 ng/µL) and were singled. mNeonGreen-positive F1 animals were scored and their development was monitored for up to five days from egg-laying. The number of fluorescent dead eggs, arrested larvae (*i*.*e*. animals never reaching adulthood) or adults were scored for each group.

### Comparison of split-sfCherry3 to split-wrmScarlet in muscle

Ten N2E wild-type animals were microinjected with either *Pmyo-3::mTagBFP2::split-wrmScarlet*_*11*_*::T2A::mNeonGreen::split-wrmScarlet*_*1-10*_*::fib-1 3 R*, or *Pmyo-3::mTagBFP2::sfCherry3*_*11*_*::T2A::mNeonGreen::sfCherry3*_*1-10*_*::fib-1 3 R* constructs were microinjected at (20 ng/µL). F_1_ animals expressing mNeonGreen in muscle were selected for comparison.

### Lifespan assays

NGM plates were supplemented with 5-Fluorouracil (5-FU, Sigma, 15 µM) (Goudeau *et al*. 2011) in order to prevent progeny from hatching and with kanamycin sulfate to prevent bacterial contamination (Sigma, 25 µg/mL). Animals fed with kanamycin-resistant OP50 were scored manually as dead or alive, from their L4 larval stage defined as day 0. A worm was considered alive if it moved spontaneously or, in cases where it wasn’t moving, if it responded to a light touch stimulus with a platinum wire. Animals that crawled off the plates, had eggs that accumulated internally, burrowed or ruptured were censored and included in the analysis until the time of censorship.

### Structure prediction and rendering of split-wrmScarlet

Phyre2 was used to predict the three-dimensional modelling in intensive mode with default parameters (Kelley *et al*. 2015). The 3D model obtained was visualized using PyMOL (v2.2.0).

### Statistical analysis

Differences in fluorescence intensity between groups were compared using unpaired *t*-test with Welch’s correction. Data are presented as means ± SD. Kaplan-Meier estimates of survival curves were calculated using *survival* (v2.38-3) and *rms* (v4.5-0) R packages and differences were tested using log-rank test. The number of animals used in each experiment is indicated in the figure legends.

### Data availability

Strains expressing a single-copy of split-wrmScarlet_1-10_ and/or sfGFP_1-10_ CF4582 (https://cgc.umn.edu/strain/CF4582), CF4587 (https://cgc.umn.edu/strain/CF4587), CF4588 (https://cgc.umn.edu/strain/CF4588), CF4610 (https://cgc.umn.edu/strain/CF4610), DUP223 (https://cgc.umn.edu/strain/DUP223) and DUP237 (https://cgc.umn.edu/strain/DUP237) are available via the *Caenorhabditis* Genetics Center (CGC). The vectors pJG100 carrying *Peft-3::split-wrmScarlet*_*1-10*_*::unc-54 3 R* (www.addgene.org/138966), pJG103 carrying split-wrmScarlet_11_ x3 tandem repeats (www.addgene.org/158533), yeast plasmids p416-TEF-membrane localization signal-mTagBFP-split-wrmScarlet_11_-TEF terminator (www.addgene.org/158585) and pRS423-GPD-split-wrmScarlet_1-10_-CYC1 terminator (www.addgene.org/158584) are deposited, along with sequences and maps at Addgene. Other strains and plasmids are available upon request. The authors state that all data necessary for confirming the conclusions presented here are represented fully within the article. A detailed protocol to generate *C. elegans* with sfGFP_11_ and/or split-wrmScarlet_11_ integrants is available at dx.doi.org/10.17504/protocols.io.bamkic4w. Supplementary material will be available at Figshare.

## RESULTS

### Split-wrmScarlet

To engineer split-wrmScarlet, first we introduced a 32 amino acid spacer between the 10^th^ and 11^th^ B-strands of full-length yeast-codon optimized mScarlet, following a strategy described previously (Feng *et al*. 2017). We subjected the spacer-inserted mScarlet sequence to several rounds of semi-random mutagenesis in *E. coli*, generating a version with fluorescence comparable to the full-length mScarlet when expressed in bacteria. However, upon separating the two fragments into two *S. cerevisiae* plasmids to test for complementation, we observed no detectable fluorescence in yeast. We decided to continue with several rounds of selection of new mutant libraries in yeast using FACS, by fusing the small fragment (without the MDELYK C-terminus residues) from our brightest *E. coli* clone to a plasma-membrane-targeted blue FP (mTagBFP), and expressing the large fragment from a high-copy number vector containing a strong promoter. The brightest resulting protein, which we named split-wrmScarlet, contained 10 amino acid substitutions relative to the C-terminal truncated mScarlet (Figure S1, A and B).

Fluorescence microscopy of yeast containing both plasmids corroborated that split-wrmScarlet showed the expected localization and can reach brightness comparable to that of intact mScarlet in yeast (Figure S2, A and B).

### Split-wrmScarlet is three-fold brighter than split-sfCherry3 in *C. elegans* muscles

In order to compare split-wrmScarlet to split-sfCherry3, the brightest published red split-FP at the time of the experiment, we combined the FP_1-10_ and FP_11_ fragments into a single plasmid for each fluorophore. Specifically, we generated worm-codon-optimized plasmids encoding three nuclear localization signals (NLS), mTagBFP2, FP_11_, a T2A peptide-bond-skipping sequence, mNeonGreen and the corresponding FP_1-10_, driven by the *eft-3* promoter (Figure S3A). Each FP_11_ was linked to mTagBFP2 in order to reduce the risk of proteolysis of the short peptide, and mNeonGreen was linked to FP_1-10_ to monitor its expression, and for normalization purposes.

Each construct was injected into wild-type animals and fluorescent progeny were analyzed. Unexpectedly, split-sfCherry3 turned out to be toxic when expressed ubiquitously, whereas 99% of split-wrmScarlet-overexpressing worms became viable adults (Figure S3B).

In an attempt to reduce split-sfCherry3-associated toxicity, we modified our construct by using the muscle-specific *myo-3* promoter and removing the NLS sequence (Figure 1B). We did not detect toxicity associated with the expression of these constructs and were able to compare the fluorescence of split-sfCherry3 and split-wrmScarlet in young adults. Red fluorescence emitted from split-wrmScarlet was 2.9-fold higher than that of split-sfCherry3 when normalized to the mNeonGreen signal (Figure 1B, 1C and S4). We also observed a 60% higher expression level of mTagBFP2 in the split-wrmScarlet-expressing animals (Supplementary Figure S4). It is worth noting that differences in expression levels could influence both brightness and toxicity comparisons. A more controlled way to compare the split FPs at similar expression levels would be to make single-copy genomic insertions of these constructs at a neutral site in the genome.

### Split-wrmScarlet_11_-mediated tagging in all somatic tissues or specifically in muscles

Our protein-tagging approach was analogous to existing split-FP methods developed for human cells (Kamiyama *et al*. 2016, Leonetti *et al*. 2016) and *C. elegans* (Hefel and Smolikove 2019). It requires split-wrmScarlet_1-10_ (*i*.*e*. just the large fragment of split-wrmScarlet without the 11th β -strand) to be expressed in the cell or tissue of interest, and the small split-wrmScarlet_11_ fragment to be inserted at an endogenous locus to tag a protein of interest (Figure 1A).

To build strains expressing single-copy insertions of split-wrmScarlet_1-10_, we first optimized its sequence for *C. elegans* codon usage (Redemann *et al*. 2011) and included three introns (Table S1). The strain expressing split-wrmScarlet_1-10_ throughout the soma (driven by the *eft-3* promoter and *unc-54* 3’UTR) was generated by editing the genome of the existing MosSCI line CA1200 (Zhang *et al*. 2015) and replacing the sequence encoding *tir-1::mRuby* with *split-wrmScarlet*_*1-10*_ using CRISPR/Cas9 and hybrid DNA templates (Paix *et al*. 2015; Dokshin *et al*. 2018; Supplementary Material, Table S4). In order to perform tissue-specific labeling, we generated a strain expressing muscle-specific split-wrmScarlet_1-10_ using the SKI-LODGE system in the strain WBM1126 (Silva-García *et al*. 2019; Supplementary Material, Table S4). The expression of split-wrmScarlet_1-10_ in these two lines did not affect the number of viable progeny (Figure S5A) nor lifespan (Figure S5B and Table S6), suggesting that the expression of split-wrmScarlet_1-10_ had no deleterious effect. To tag a gene of interest with the split-wrmScarlet_11_ fragment, we used microinjection of preassembled Cas9 ribonucleoproteins, because this method enables high-efficiency genome editing in worms (Paix *et al*. 2015). The most efficient insertion of short sequences in *C. elegans* was previously shown to be achieved using ssODN donors (Paix *et al*. 2015; Prior *et al*. 2017; Dokshin *et al*. 2018). A great advantage of this strategy is that all of the components required for editing are commercially available or can be synthesized rapidly in the lab (Leonetti *et al*. 2016). Synthetic ssODNs have a typical size limit of 200 nt. The small size of split-wrmScarlet_11_ (18-24 a.a.) is key: 200 nt can encompass split-wrmScarlet_11_ (66-84 nt, including a 4 a.a. linker) flanked by two homology arms > 34 nt (up to 67-58 nt) for HDR. In principle, a few days after the somatic and/or muscle-specific split-wrmScarlet_1-10_ strain(s) are microinjected, progeny can be screened for red fluorescence, genotyped and sequenced to check the accuracy of editing (Figure 2; a detailed protocol is available at dx.doi.org/10.17504/protocols.io.bamkic4w). If desired, co-CRISPR strategies such as *dpy-10*(*cn64*) (Paix *et al*. 2015) or co-injection with pRF4 (Dokshin *et al*. 2018) can be used to screen for correct candidates and to control for microinjection efficacy and payload toxicity.

**Figure 2.**
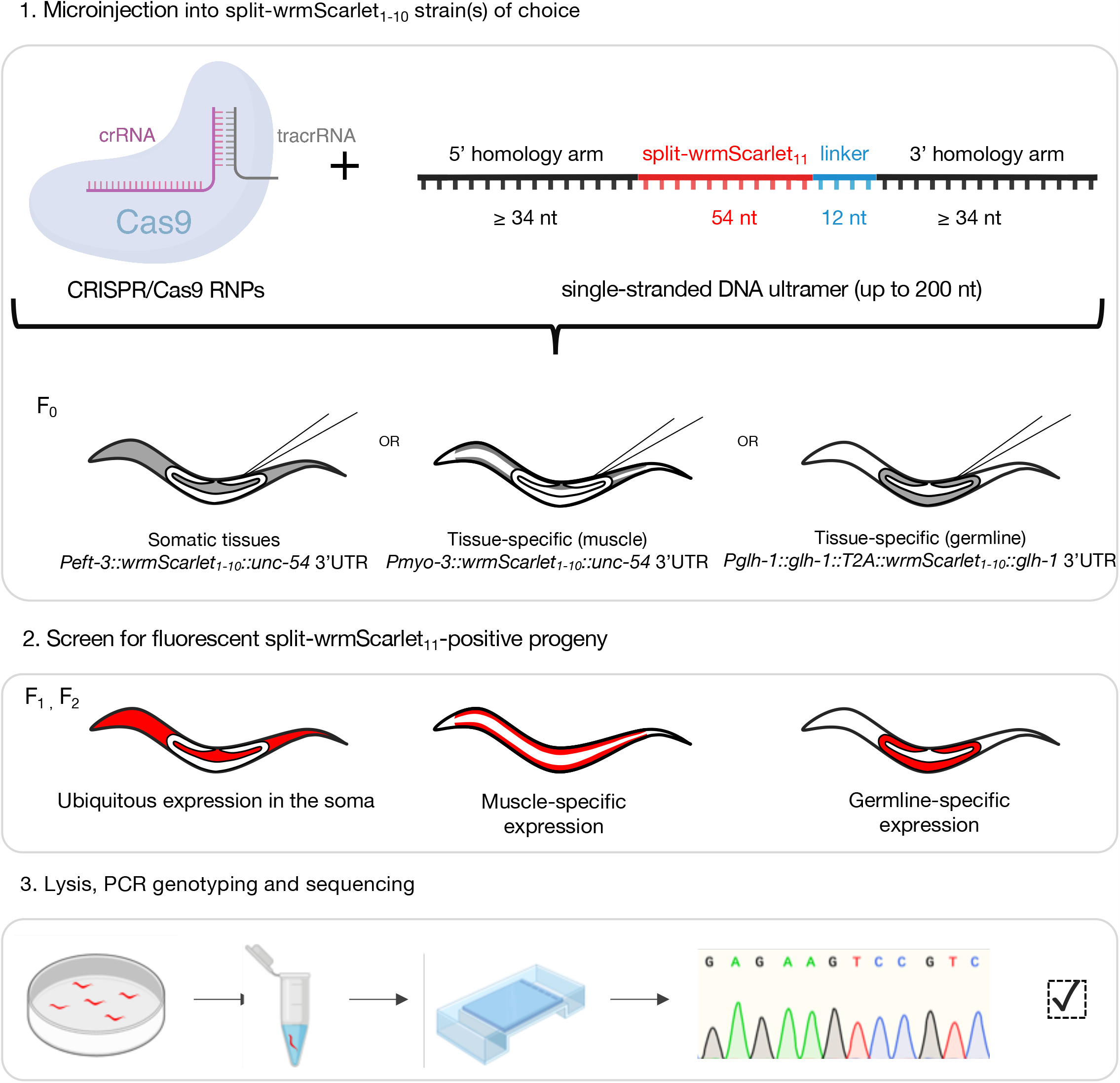
Split-wrmScarlet_11_-mediated tagging. Schematic representation of the split-wrmScarlet workflow to visualize endogenous proteins specifically in muscles, germline, or throughout the soma. Some illustrations were created with BioRender.com.

To test our approach, we used it to tag six proteins with distinct subcellular localizations. Starting with the somatic split-wrmScarlet_1-10_ parental strain CF4582, we introduced split-wrmScarlet_11_ at the N-terminus of TBB-2, FIB-1 or VHA-13 or at the C-terminus of EAT-6, HIS-3 and TOMM-20 (Supplementary Material, Table S4). These proteins mark the cytoskeleton, nucleoli, lysosomes, plasma membrane, nuclei and mitochondria, respectively. Importantly, for tagging transmembrane proteins, the split-wrmScarlet_11_ tag was introduced at the terminus exposed to the cytosol. Split-wrmScarlet fluorescence from all six proteins matched their expected subcellular localization in somatic cells (Figure 3, A-F). To test the muscle-specific split-wrmScarlet_1-10_ line CF4610, we tagged the N-terminus of endogenous FIB-1 with split-wrmScarlet_11_ and confirmed the fluorescence from nucleoli in muscle cells (Figure 3G). Together, our results show that split-wrmScarlet enables rapid fluorescent tagging of proteins with disparate cytoplasmic or nuclear locations expressed from their endogenous loci.

**Figure 3.**
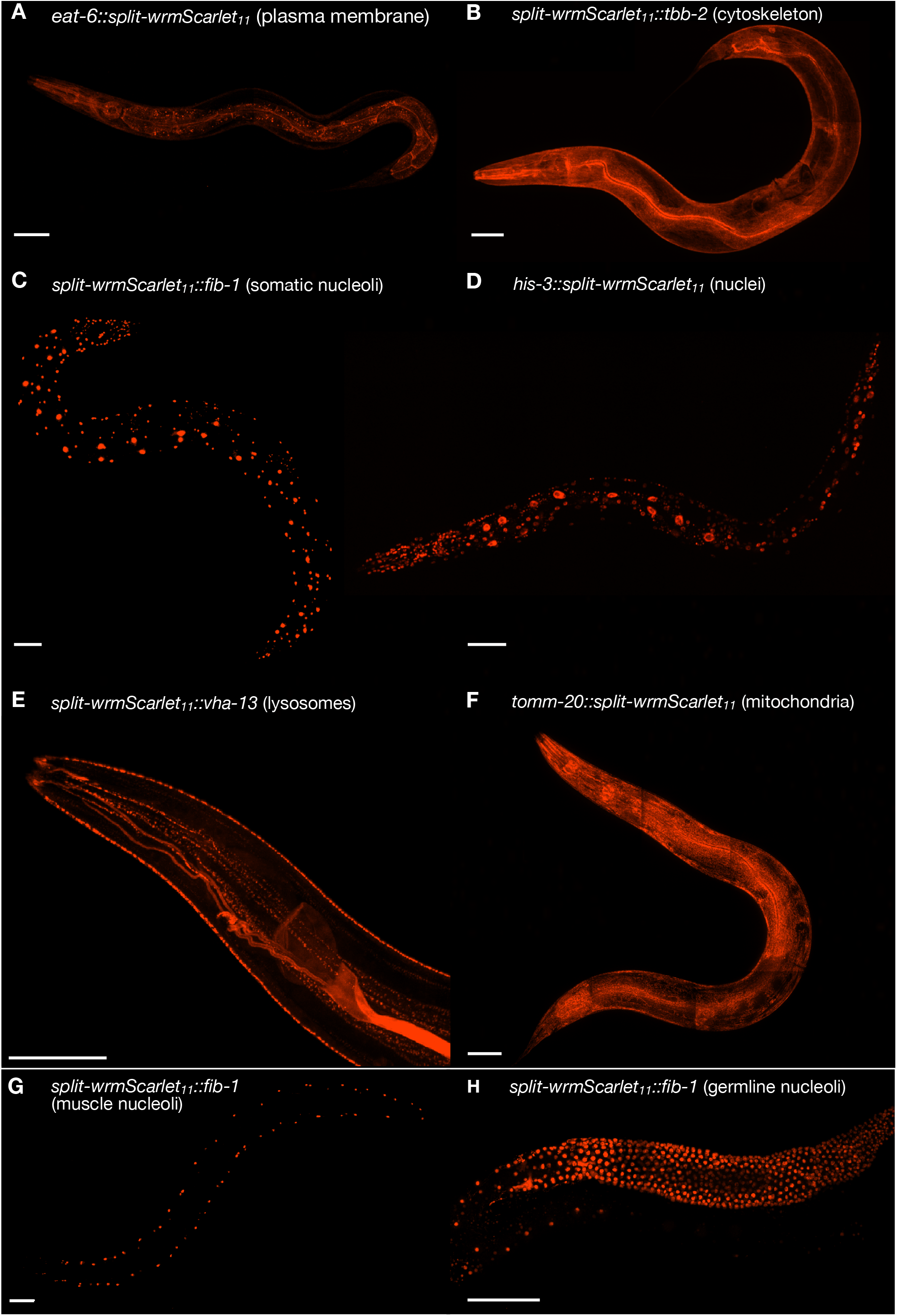
Split-wrmScarlet labeling of proteins with distinct subcellular locations. Endogenous proteins tagged with split-wrmScarlet_11_ in animals expressing split-wrmScarlet_1-10_ in somatic tissues, in muscles or in the germline. (A-F) Confocal images of worms expressing somatic split-wrmScarlet_1-10_ and (A) EAT-6::split-wrmScarlet_11_ (plasma membrane), (B) split-wrmScarlet_11_::TBB-2 (cytoskeleton), (C) split-wrmScarlet_11_::FIB-1 (nucleoli), (D) HIS-3::split-wrmScarlet_11_ (nuclei), (E) split-wrmScarlet_11_::VHA-13 (lysosomes), or (F) TOMM-20::split-wrmScarlet_11_ (mitochondria). (G) Transgenic worm expressing split-wrmScarlet_1-10_ in muscle and split-wrmScarlet_11_::FIB-1. (H) Transgenic worm expressing split-wrmScarlet_1-10_ in the germline and split-wrmScarlet_11_::FIB-1. (A-H) Maximum intensity projections of 3D stacks shown. Scale bars, 50 µm.

The 18 a.a. split-wrmScarlet_11_ sequence used for these experiments ends with two glycines. In mammalian cells, C-terminal gly-gly sequences have been reported to function as degron signals (Koren *et al*. 2018). Our TOMM-20::split-wrmScarlet_11_ had a spontaneous mutation of the last glycine to a stop codon (Supplementary Material, Table S4), which could be problematic if the protein degradation mechanism, DesCEND (destruction via C-end degrons) operates in *C. elegans*. However, we do not detect differences in protein abundance of HIS-3 vs HIS-3:split-wrmScarlet_11_ by western blots in *C. elegans* (Figure S11). We also do not detect differences in protein abundance of mScarlet truncated to end in gly-gly compared to mScarlet ending in MDELYK via fluorescence in yeast. Nonetheless, we recommend using a 24 a.a. split-wrmScarlet_11_ sequence YTVVEQYEKSVARHCTGGMDELYK when labeling proteins at their C-terminus to avoid the possibility that split-wrmScarlet_11_ ending in gly-gly could function as a degron. This modified sequence still fits within the 200 nt ssODN synthesis limit and works at least as well as the 18 a.a. split-wrmScarlet_11_ sequence (Figure S6).

### Split-wrmScarlet_11_-mediated tagging in the germline

Our initial attempt to use split-wrmScarlet in the germline failed. We made a single-copy integrated *Psun-1::split-wrmScarlet*_*1-10*_*::sun-1* 3’UTR strain via MosSCI, but when we injected a plasmid encoding mNeonGreen::split-wrmScarlet_11_, we observed green fluorescence, but no red fluorescence (Figure S7, A and B), suggesting the absence of split-wrmScarlet_1-10_ expression. We suspected germline silencing of the germline-expressed *split-wrmScarlet*_*1-10*_, so we attempted an alternative expression approach which we call “glonads”. The germline-helicase protein GLH-1 is highly expressed and germline-specific (Marnik *et al*. 2019). We fused a *T2A::split-wrmScarlet*_*1-10*_ sequence to the C-terminus of the endogenous *glh-1* gene using CRISPR/Cas9. The high expression of GLH-1 yielded high expression of split-wrmScarlet_1-10_, and the T2A separated split-wrmScarlet_1-10_ from GLH-1 (Liu *et al*. 2017). The *glh-1::T2A::split-wrmScarlet*_*1-10*_ strain (https://cgc.umn.edu/strain/DUP237) can be used like our other tissue-specific strains for germline-specific tagging. To demonstrate this, we tagged the N-terminus of endogenous FIB-1 with split-wrmScarlet_11_, and we observed red fluorescence localized to the nucleoli specifically in the germline and embryos, as we hoped (Figure 3H; Figure S8, A and C). Finally, we note that this *glh-1::T2A::X* strategy is not limited to split-wrmScarlet; where *X* could potentially be any fluorescent protein, or more generally, any exogenous protein one wish to express in a germline-specific way using CRISPR/Cas9.

### Split-wrmScarlet_11_ tandem repeats increase fluorescence

To benchmark the fluorescence intensity of split-wrmScarlet against its full-length counterpart, we first generated *wrmScarlet::vha-13* (El Mouridi *et al*. 2017) transgenic animals and compared their fluorescence to *split-wrmScarlet*_*11*_*::vha-13* in worms expressing split-wrmScarlet_1-10_ somatically (Figure 4, A and B). At the *vha-13* locus, split-wrmScarlet was about half as bright as a full-length fluorophore (48%), a ratio comparable to that of split-mNeonGreen2 and its full-length counterpart in human cells (Feng *et al*. 2017).

**Figure 4.**
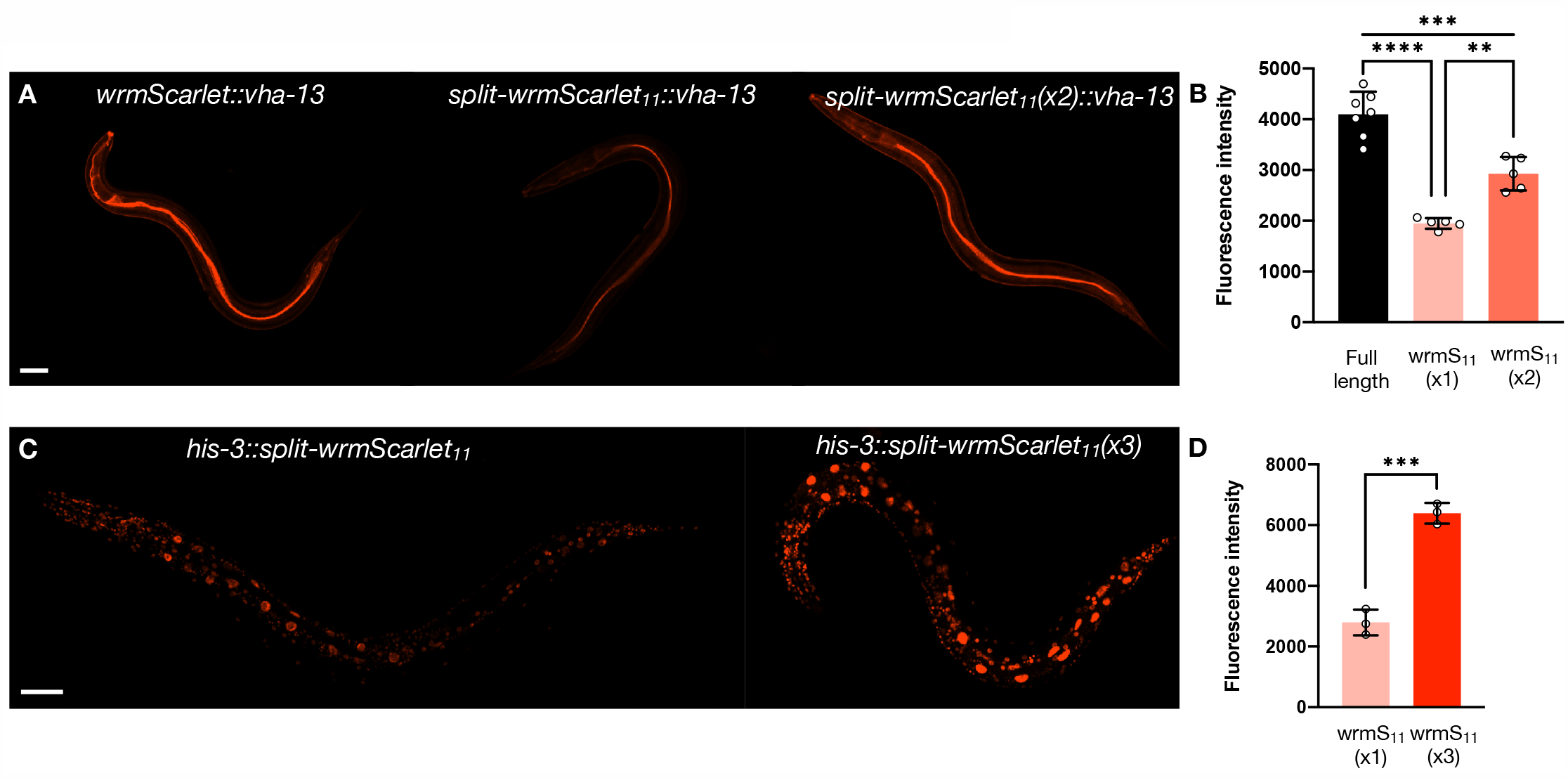
Split-wrmScarlet_11_ tandem repeats increase fluorescence. (A) Images of animals carrying either wrmScarlet, split-wrmScarlet_11_ or two tandem repeats of split-wrmScarlet_11_ inserted at the endogenous VHA-13 N-terminus. (B) Emission intensities of animals carrying wrmScarlet, split-wrmScarlet_11_ or dual split-wrmScarlet_11_ inserted at the VHA-13 N-terminus. Mean ± s.d. Circles are individuals. \*\*\*\**P* < 0.0001, \*\*\**P* < 0.001, \*\**P* < 0.005. (C) Images of animals carrying either a single split-wrmScarlet_11_ or three tandem repeats of split-wrmScarlet_11_ inserted at the HIS-3 C-terminus. (D) split-wrmScarlet emission intensities from animals carrying a single split-wrmScarlet_11_ or three tandem repeats of split-wrmScarlet_11_ knock-in at the HIS-3 C-terminus. Mean ± s.d. Circles are individuals. \*\*\**P* < 0.001. Images from each comparison were taken under identical instrument conditions using confocal microscopy and are shown using identical brightness and contrast settings. Images shown are from a single confocal plane. Scale bars, 50 µm.

Since visualizing endogenous proteins of low abundance can be challenging, it is key to address this limitation. Increasing the number of FP_11_ domains tagged to an endogenous protein multiplies the number of the corresponding FP_1-10_s recruited, increasing the overall fluorescent signal in human cells (Leonetti *et al*. 2016) and in *C. elegans* (He *et al*. 2019; Hefel and Smolikove 2019). To demonstrate that split-wrmScarlet fluorescence is enhanced by split-wrmScarlet_11_ tandem repeats, we introduced two split-wrmScarlet_11_ domains at the N-terminus of *vha-13* and three split-wrmScarlet_11_ domains at the C-terminus of *his-3* in animals expressing somatic split-wrmScarlet_1-10_. Compared to animals carrying a single split-wrmScarlet_11_ at the identical locus, carrying two split-wrmScarlet_11_s increased overall fluorescence by 1.5-fold, while carrying three increased it by 2.3-fold (Figure 4, C and D). Note that our three-split-wrmScarlet_11_ tandem sequence exceeds the 200 nt ssODN synthesis limit, so we used dsDNA donor templates for these constructions (Supplementary Material, Table S4).

### sfGFP_11_-mediated tagging in somatic cells

Split-sfGFP has been used successfully in worms before (Noma *et al*. 2017; He *et al*. 2019; Hefel and Smolikove 2019). However, there is still a need for a strain that ubiquitously expresses sfGFP_1-10_ in the soma from an integrated single-copy insertion in order to avoid heterogeneous expression and time-consuming manual maintenance. To build this strain, we codon-optimized the original sfGFP_1-10_ sequence for *C. elegans* and included one intron (Cabantous *et al*. 2004; Redemann *et al*. 2011; Supplementary Material, Table S1). We initially generated a strain expressing sfGFP_1-10_ driven by the *let-858* promoter and *unc-54* 3’UTR using MosSCI, but later replaced the *let-858* promoter with the *eft-3* promoter using CRISPR/Cas9 and hybrid DNA donor template because we observed that *Peft-3* resulted in significantly higher levels of gene expression (Paix *et al*. 2015; Dokshin *et al*. 2018; Supplementary Material, Table S4). To validate this strain, we inserted sfGFP_11_ at the N-terminus of lysosomal VHA-13 or at the C-terminus of nuclear-localized HIS-3 (Figure 5, A and B). Both strains yielded relatively bright signals in accordance with their predicted subcellular localization. We generated eGFP::VHA-13 transgenic animals and compared their fluorescence to sfGFP_11_::VHA-13 in worms expressing sfGFP_1-10_ somatically (Figure S9, A and B). At the *vha-13* locus, split-sfGFP was about a third as bright as a full-length eGFP. It is worth noting that this comparison is not perfect, in part due to the absence of the six superfolder mutations S30R, Y39N, N105T, Y145F, I171V and A206V in the eGFP fluorophore.

**Figure 5.**
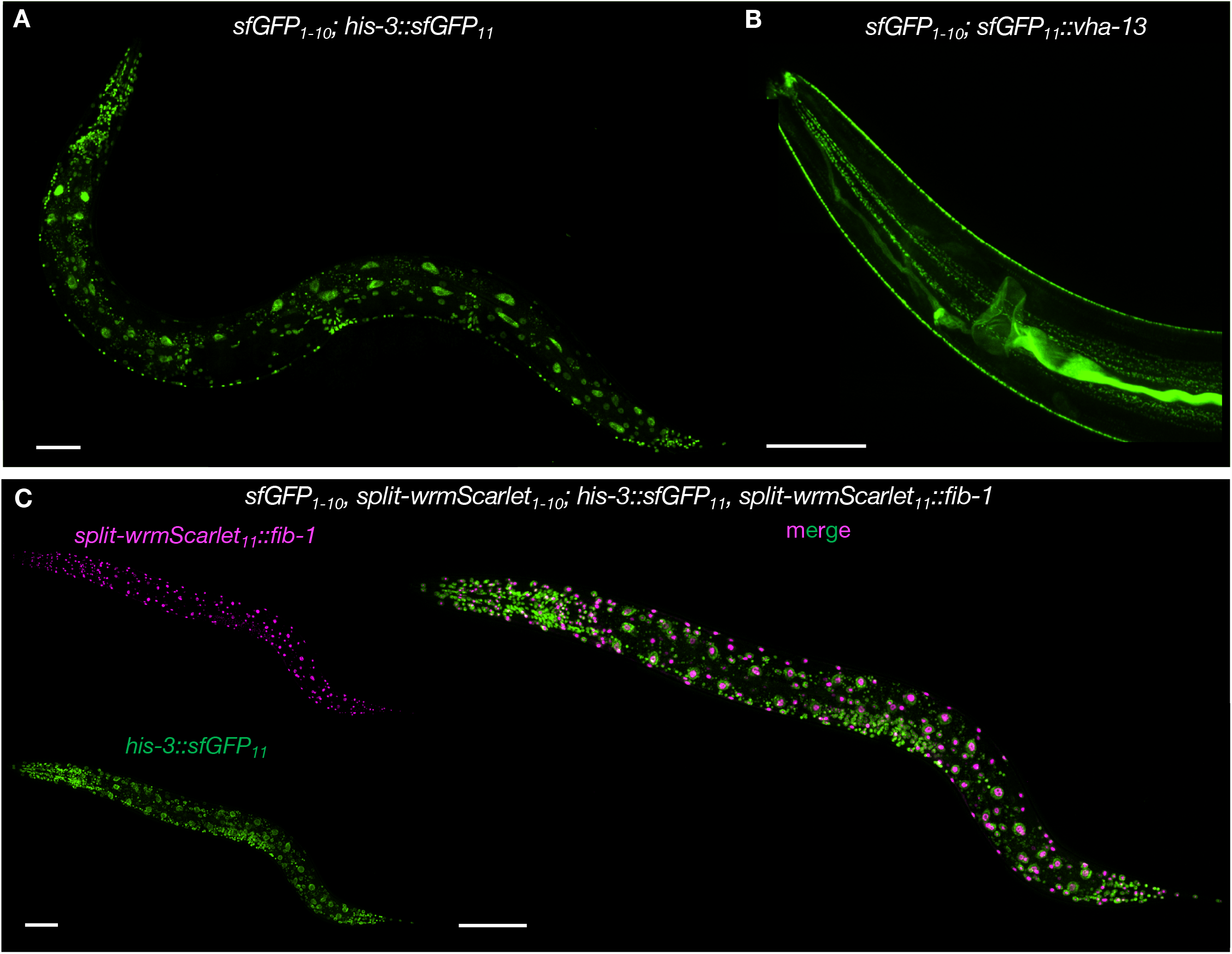
Split-sfGFP and split-wrmScarlet dual-color protein labeling. Images of animals stably expressing sfGFP_1-10_ in somatic tissues (A) CF4592 (*muIs253*[*Peft-3::sfGFP*_*1-10*_*::unc-54 3’UTR, Cbr-unc-119*(*+*)] *II; unc-119*(*ed3*) *III; his-3*(*muIs255*[*his-3::sfGFP*_*11*_] *V*) or (B) CF4589 (*muIs253*[*Peft-3::sfGFP*_*1-10*_*::unc-54 3’UTR, Cbr-unc-119*(*+*)] *II; unc-119*(*ed3*) *III; vha-13*(*muIs268*[*sfGFP*_*11*_*::vha-13*]) *V*). (C) Dual color protein labeling with split-wrmScarlet and split-sfGFP in somatic cells. Composite display of red and green channels of animals expressing split-wrmScarlet_1-10_ and sfGFP_1-10_ in somatic tissues, HIS-3::sfGFP_11_ and split-wrmScarlet_11_::FIB-1; CF4602 (*muIs253*[*Peft-3::sfGFP*_*1-10*_*::unc-54 3’UTR, Cbr-unc-119*(*+*)], *muIs252*[*Peft-3::split-wrmScarlet*_*1-10*_*::unc-54 3’UTR, Cbr-unc-119*(*+*)] *II; unc-119*(*ed3*) *III; fib-1*(*muIs254*[*split-wrmScarlet*_*11*_*::fib-1*]), *his-3*(*muIs255*[*his-3::sfGFP*_*11*_]) *V*). Maximum intensity projections of 3D stacks shown. Scale bars, 50 µm.

### sGFP2_11_-mediated tagging in the germline

We also generated a germline-specific sGFP2_1-10_ strain using a similar strategy (Figure S8B, Supplementary Material, Table S1). Split-sGFP2 is a split-superfolder GFP variant optimized for brightness and photostability (Köker *et al*. 2018). To test this germline-specific line DUP223 (https://cgc.umn.edu/strain/DUP223), we tagged the C-terminus of endogenous PGL-1 with sGFP2_11_ and confirmed the green fluorescence from P-granules (Figure S8B, lower panel).

### Dual color protein labeling with split-wrmScarlet and split-sfGFP

Finally, to allow two-color imaging in the soma, we crossed the strains *Peft-3::sfGFP*_*1-10*_; *his-3::sfGFP*_*11*_ (CF4592) and *Peft-3::split-wrmScarlet*_*1-10*_; *split-wrmScarlet*_*11*_*::fib-1* (CF4601). This cross resulted in the line *Peft-3::sfGFP*_*1-10*_, *Peft-3::split-wrmScarlet*_*1-10*_ (https://cgc.umn.edu/strain/CF4588) as well as the dually labeled strain *Peft-3::sfGFP*_*1-10*_, *Peft-3::split-wrmScarlet*_*1-10*_; *split-wrmScarlet*_*11*_*::fib-1, his-3::sfGFP*_*11*_ (CF4602, Figure 5C). The fluorescent signals from both split-FPs appeared in their respective subcellular compartments, suggesting the two systems are compatible. We note an additional advantage of the strain CF4588: the loci of *split-wrmScarlet*_*1-10*_ and *sfGFP*_*1-10*_ are genetically linked (only 0.96 cM apart), which facilitates outcrossing when needed. In addition, all our *C. elegans* lines expressing *split-wrmScarlet*_*1-10*_ and *sfGFP*_*1-10*_ are viable homozygotes, so the strains do not require special maintenance.

### The current split-wrmScarlet is not detectable in mammalian cells

We failed to detect split-wrmScarlet in mammalian cells, despite our efforts to rescue its fluorescence by screening a mammalian-codon-optimized split-wrmScarlet_11_ single/double mutant library in HEK293T cells (Figure S10, A and B, and supplementary text).

## DISCUSSION

In this study, we describe several new tools for rapid CRISPR-mediated labeling of endogenously-expressed proteins using split fluorophores. While these tools are powerful and relatively easy to implement, several considerations should be taken into account when using this method.

First, detection of a given protein labeled with an FP_11_ can only occur in a cellular compartment where the corresponding FP_1-10_ is present. Proteins tagged with split-wrmScarlet_11_ or sfGFP_11_ generated in this work were either exposed to the cytosol or nucleoplasm (nuclei or nucleoli), where split-wrmScarlet_1-10_ and/or sfGFP_1-10_ were present. For proteins or epitopes located within the lumen of organelles, such as mitochondria or the endoplasmic reticulum, one might need to generate and validate *C. elegans* lines expressing split-wrmScarlet_1-10_ or sfGFP_1-10_ containing a mitochondrial localization sequence or ER signal peptide and retention signals, respectively. These approaches have been used successfully in mammalian cells with split-sfGFP when tagging ER-resident polypeptides (Kamiyama *et al*. 2016) and with split-sfCherry2 to detect protein present in the mitochondrial matrix (Ramadani-Muja *et al*. 2019).

Second, when labeling proteins with split-wrmScarlet at the C-terminus, we recommend using the 24 a.a. split-wrmScarlet_11_ sequence YTVVEQYEKSVARHCTGGMDELYK. As described in the Results section, our 18 a.a. split-wrmScarlet_11_ fragment ends in gly-gly, which has been shown to be a degron signal in mammalian cells. We cannot exclude the possibility that ending in gly-gly can be detrimental in *C. elegans*. The 24 a.a. split-wrmScarlet_11_ still fits within a 200 nt ssODN donor template with a 12 nt linker and up to 58 nt homology arms and is at least as bright as the 18 a.a. split-wrmScarlet_11_ (Figure S6).

Third, as for any other protein tag, it is important to select, when possible, a site that is unlikely to interfere with protein folding, function or localization (Snapp 2005; Nance and Frøkjær-Jensen 2019). For example, N-termini of membrane- and organelle-resident proteins often contain signal peptides or localization signals, and C-termini may contain degron sequences that regulate protein turnover. Interestingly, there are examples of proteins that become toxic when tagged with a full-length GFP, but tolerate labeling with a split fluorescent protein. For example, SYP-4 was reported to be mostly functional when endogenously-tagged with sfGFP_11_ in a strain expressing sfGFP_1-10_ specifically in the germline, but not functional when labeled with full-length GFP (Hefel and Smolikove 2019).

Fourth, for proteins of interest present at low levels, we provided an alternative protocol to insert an additional two or three split-wrmScarlet_11_ fragments, which increases the overall fluorescence substantially. However, the number of split-wrmScarlet_11_ fragments could likely be increased further, to at least seven tandem repeats, based on approaches used successfully with split-sfGFP in human cells (Feng *et al*. 2017) and *C. elegans* (Noma *et al*. 2017; He *et al*. 2019; Hefel and Smolikove 2019).

Fifth, we would like to emphasize differences between our technique and the bimolecular fluorescence-complementation (BiFC) assay. When used together, high-affinity green and red split fluorescent proteins can provide information on co-localization, but unlike BiFC split proteins (Hu *et al*. 2002), they are not intended to assess protein-protein interactions directly.

This is because BiFC split proteins require finely tuned weak affinities that do not disrupt the underlying interaction being studied. In our approach, only the split-wrmScarlet_11_ fragment is attached to a protein of interest, the split-wrmScarlet_1-10_ fragment is expressed in excess and unattached.

Finally, we would like to note that despite being three times brighter than the latest split-sfCherry3 in worms, our current split-wrmScarlet was not visible in the mammalian cell line we examined (Figure S10). Its ability to fluoresce is not restricted to worms, because it can reach wild-type levels of brightness in yeast. We do not know the basis for this discrepancy. It is possible that the concentration of the split-wrmScarlet_1-10_ fragment in mammalian cells is too low to drive complementation with split-wrmScarlet_11_. This could potentially be overcome by further mutagenizing split-wrmScarlet and screening for fluorescence at low expression levels in mammalian cells.

We believe our system can substantially increase the speed, efficiency, and ease of *in vivo* microscopy studies in *C. elegans*. We expect it to facilitate two-color and co-localization experiments and to find wide use in the worm community. We believe that these strains could facilitate novel or large-scale experiments, such as efforts to tag the entire genome of *C. elegans*.

## Author contributions

M.I. developed split-wrmScarlet in A.G.Y. laboratory. J.P. performed the cell sorting. J.G. performed *C. elegans* experiments in C.K. laboratory. C.S. generated the two *C. elegans* germline strains in D.U. laboratory. L.S. conducted the mammalian cell experiments in M.D.L. laboratory. J.G. wrote the initial draft. All authors provided intellectual contributions to the collaboration.

## Acknowledgements

We thank Katie Podshivalova, Rex Kerr, Calvin Jan and David Botstein for comments on the manuscript, Peichuan Zhang for experimental suggestions, Vikram Narayan for biochemistry advice, and members of the Kenyon lab for fruitful discussions. We would like to thank Abby Dernburg for bringing to our attention that a gly-gly C-terminus might be a degron. We also thank Behnom Farboud from Barbara Meyer’s laboratory for discussing CRISPR/Cas9 protocols and Liangyu Zhang from Abby Dernburg’s laboratory for sharing sequences of the CA1200 strain. Some strains were provided by the CGC, which is funded by NIH Office of Research Infrastructure Programs (P40 OD010440). This project was supported by Calico Life Sciences LLC.

## FIGURE LEGENDS

**Figure S1.**
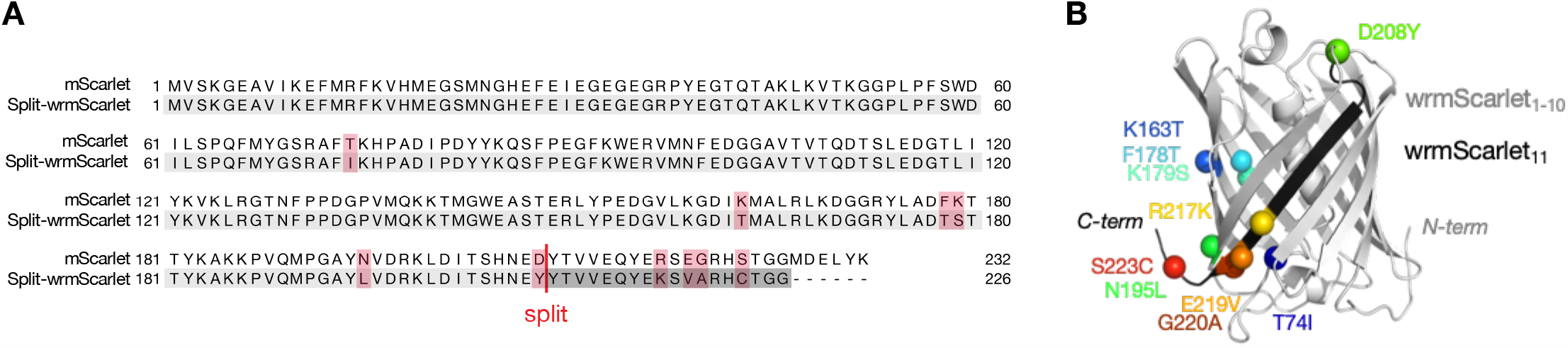
Split-wrmScarlet sequence comparison to mScarlet. (A) Protein sequence alignment of mScarlet and split-wrmScarlet. The ten amino acid substitutions are highlighted in red, the sequence of split-wrmScarlet_1-10_ in light gray and the sequence of split-wrmScarlet_11_ in dark gray. Note that gly-gly sequences at the C-termini of proteins can be degrons in mammalian cells, so when labeling on the C-terminus, we recommend using the 24 a.a. split-wrmScarlet_11(MDELYK)._ (B) Protein structure from split-wrmScarlet generated with Phyre2 and PyMOL. Mutations of split-wrmScarlet relative to mScarlet are highlighted with surface rendering, and the split-wrmScarlet_11_ strand is colored in black.

**Figure S2.**
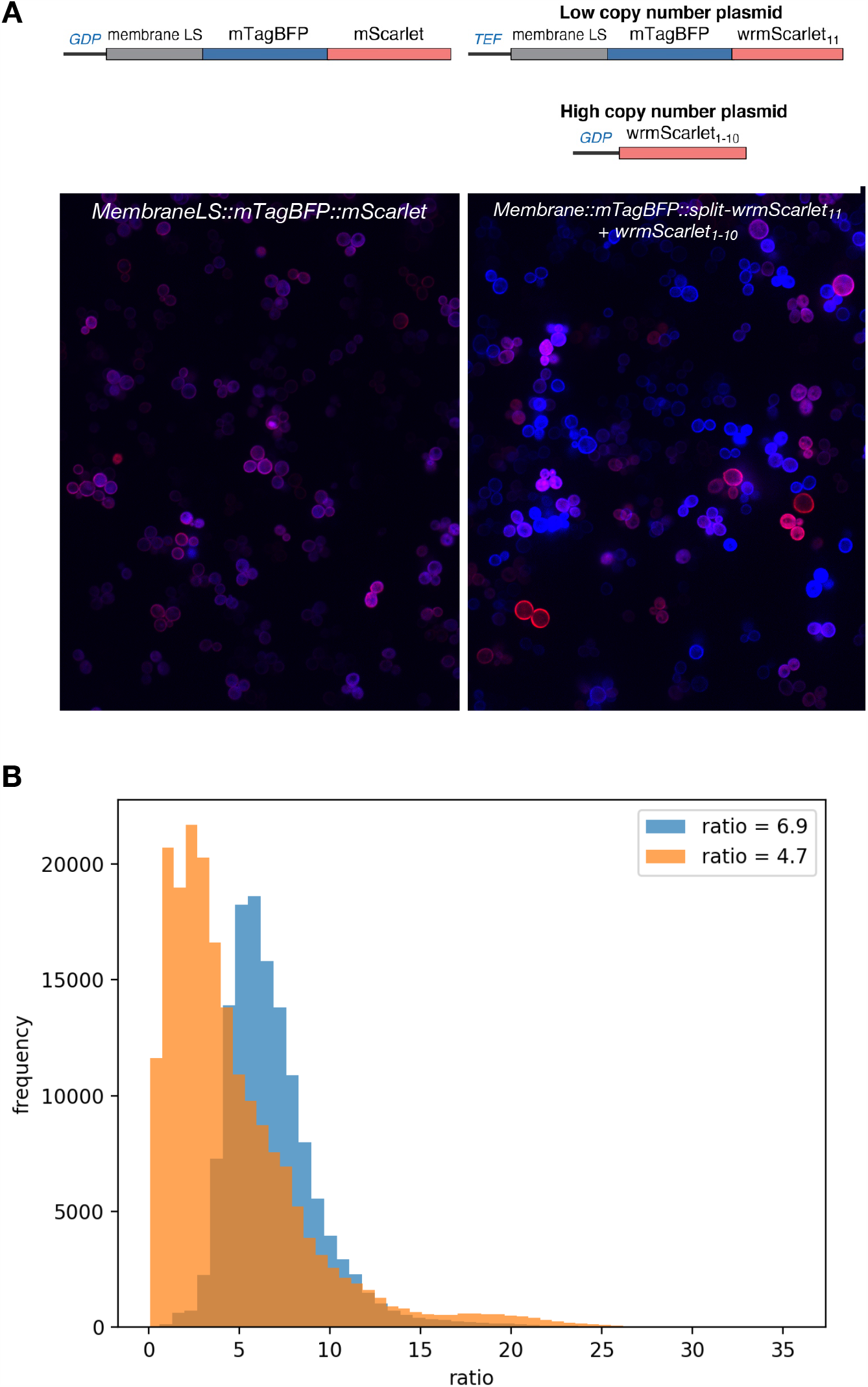
Split-wrmScarlet brightness in *S. cerevisiae*. (A) Composite display of red and blue channels for membrane-localized mTagBFP-mScarlet (wild-type) fusion or split-wrmScarlet_1-10_ plus membrane localized mTagBFP-split-wrmScarlet_11_ in yeast. Images were acquired and are displayed under identical conditions. Note that the heterogeneity inherent to expression from plasmids is large, but split-wrmScarlet is capable of brightness levels similar to the parent protein. A schematic of the plasmids transformed is presented above each image. (B) Histograms displaying the per-pixel ratio of red to blue fluorescence for background corrected, masked images. mScarlet/mTagBFP ratios are displayed in blue, and split-wrmScarlet/mTagBFP in orange. The inset displays the average red/blue ratio.

**Figure S3.**
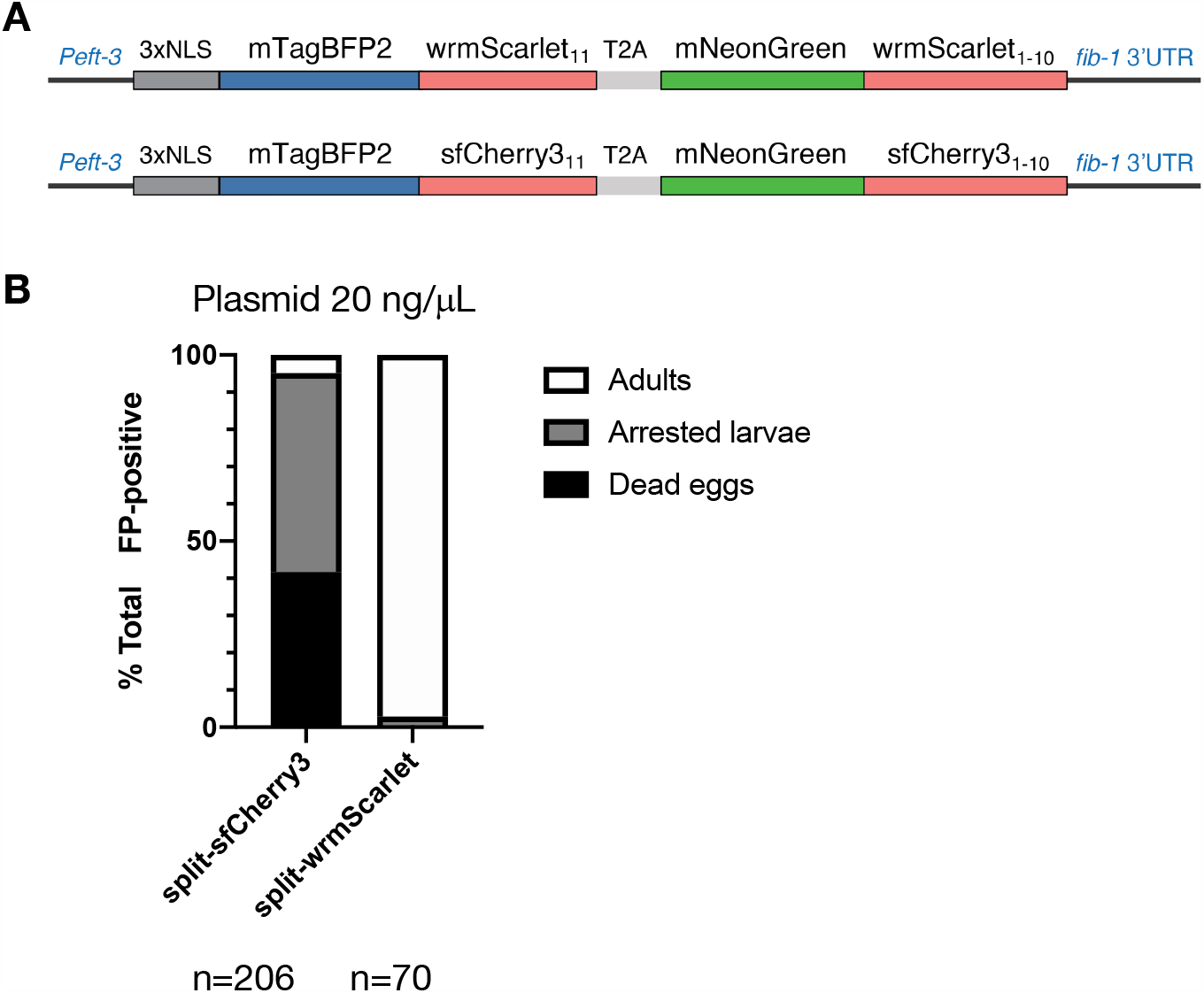
Developmental toxicity in worms expressing split-sfCherry3 in somatic nuclei. (A) Schematic of the plasmids encoding split-wrmScarlet and split-sfCherry3 used for comparison. Each plasmid consists of a large FP_1-10_ fused to mNeonGreen, and the corresponding small FP_11_ fused to mTagBFP2 preceded with three SV40 nuclear localization sequences (NLS). The T2A sequence ensures the separation of NLS::mTagBFP2::FP_11_ and the corresponding mNeonGreen::FP_1-10_. (B) Quantification of mNeonGreen-positive animals into one of three classes: dead eggs, larvae or adults.

**Figure S4.**
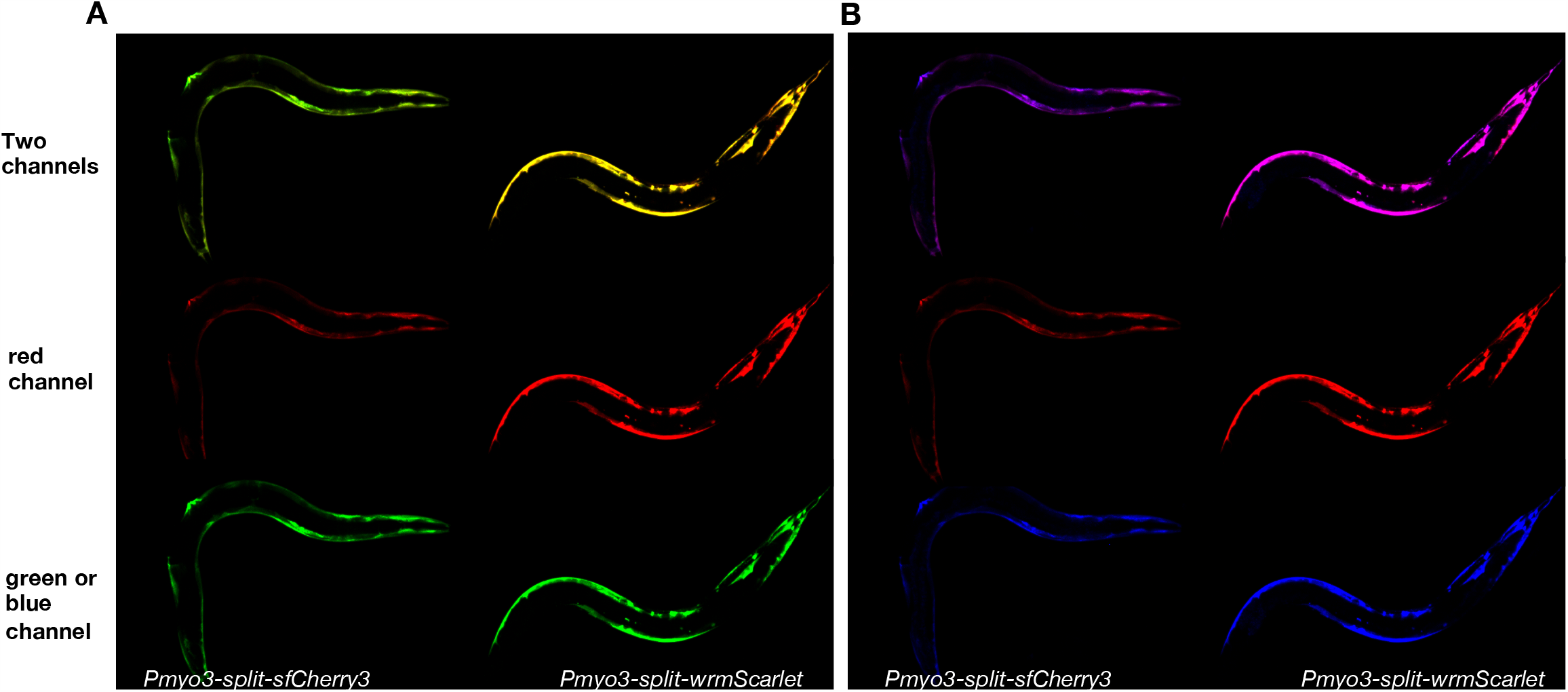
Split-wrmScarlet and split-sfCherry3 comparison. Individual channels and overlays corresponding to the images displayed in Figure 1B.

**Figure S5.**
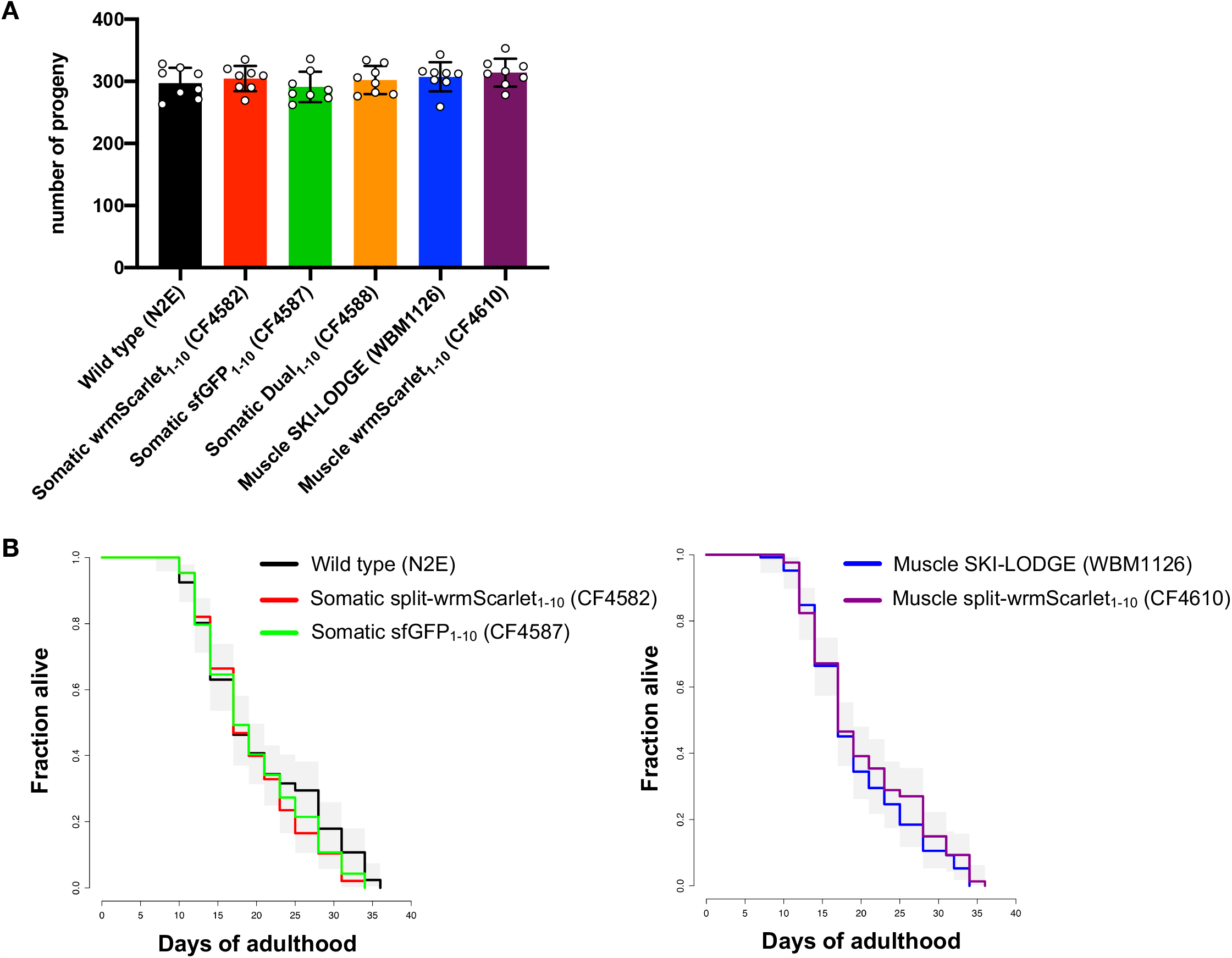
Brood size and lifespan of split-wrmScarlet_1-10_ and sfGFP_1-10_ lines. Split-wrmScarlet_1-10_ and sfGFP_1-10_ lines produced wild-type numbers of progeny (A) and a wild-type lifespan (B). Genotypes: N2E (wild-type), CF4582 (*muIs252*[*Peft-3::split-wrmScarlet*_*1-10*_*::unc-54 3’UTR, Cbr-unc-119*(*+*)] *II; unc-119*(*ed3*) *III*), CF4587 (*muIs253*[(*Peft-3::sfGFP*_*1-10*_*::unc-54 3’UTR, Cbr-unc-119*(*+*)] *II; unc-119*(*ed3*) *III*), CF4588 (*muIs253*[*Peft-3::sfGFP*_*1-10*_*::unc-54 3’UTR, Cbr-unc-119*(*+*)], *muIs252*[*Peft-3::split-wrmScarlet*_*1-10*_*::unc-54 3’UTR, Cbr-unc-119*(*+*)] *II; unc-119*(*ed3*) *III*), CF4610 (*muIs257*[*Pmyo-3::split-wrmScarlet*_*1-10*_*::unc-54 3’UTR*] *I*) and WBM1126 (*wbmIs61*[*myo-3p::3XFLAG::dpy-10 crRNA::unc-54 3’UTR*] *I*). Supplementary Material, Table S6 shows survival statistics for all lifespan experiments.

**Figure S6.**
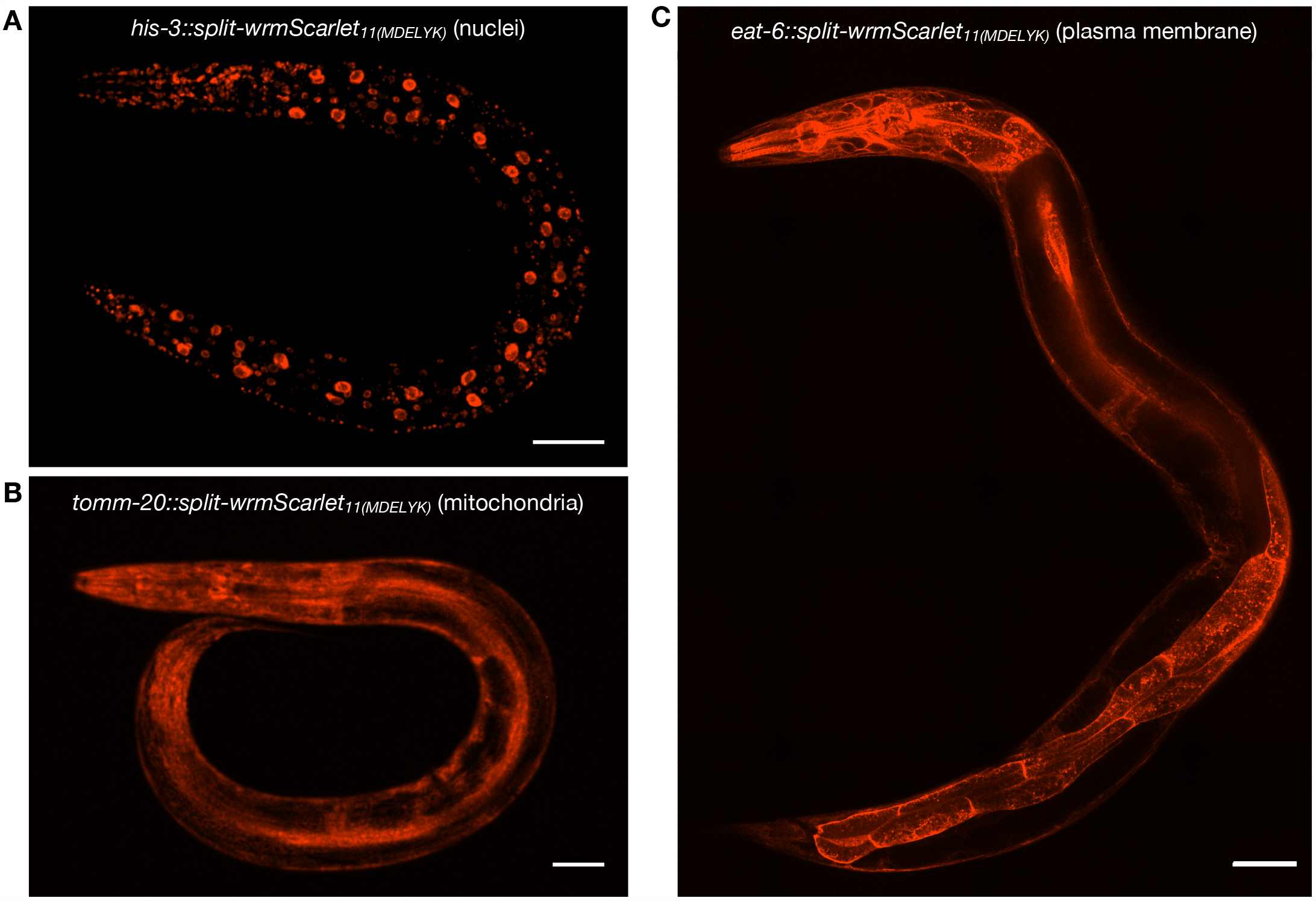
Proteins tagged at their C-terminus with the 24 a.a. split-wrmScarlet_11(MDELYK)_ Endogenous proteins tagged with split-wrmScarlet_11(MDELYK)_ in animals expressing split-wrmScarlet_1-10_ in somatic tissues. (A-C) Confocal images of worms expressing somatic split-wrmScarlet_1-10_ and (A) HIS-3::split-wrmScarlet_11(MDELYK)_ (nuclei), (B) TOMM-20::split-wrmScarlet_11(MDELYK)_ (mitochondria), or (C) EAT-6::split-wrmScarlet_11(MDELYK)_ (plasma membrane). (A-B) Maximum intensity projections of 3D stacks shown. (C) Single slice shown. Scale bars, 50 µm.

**Figure S7.**
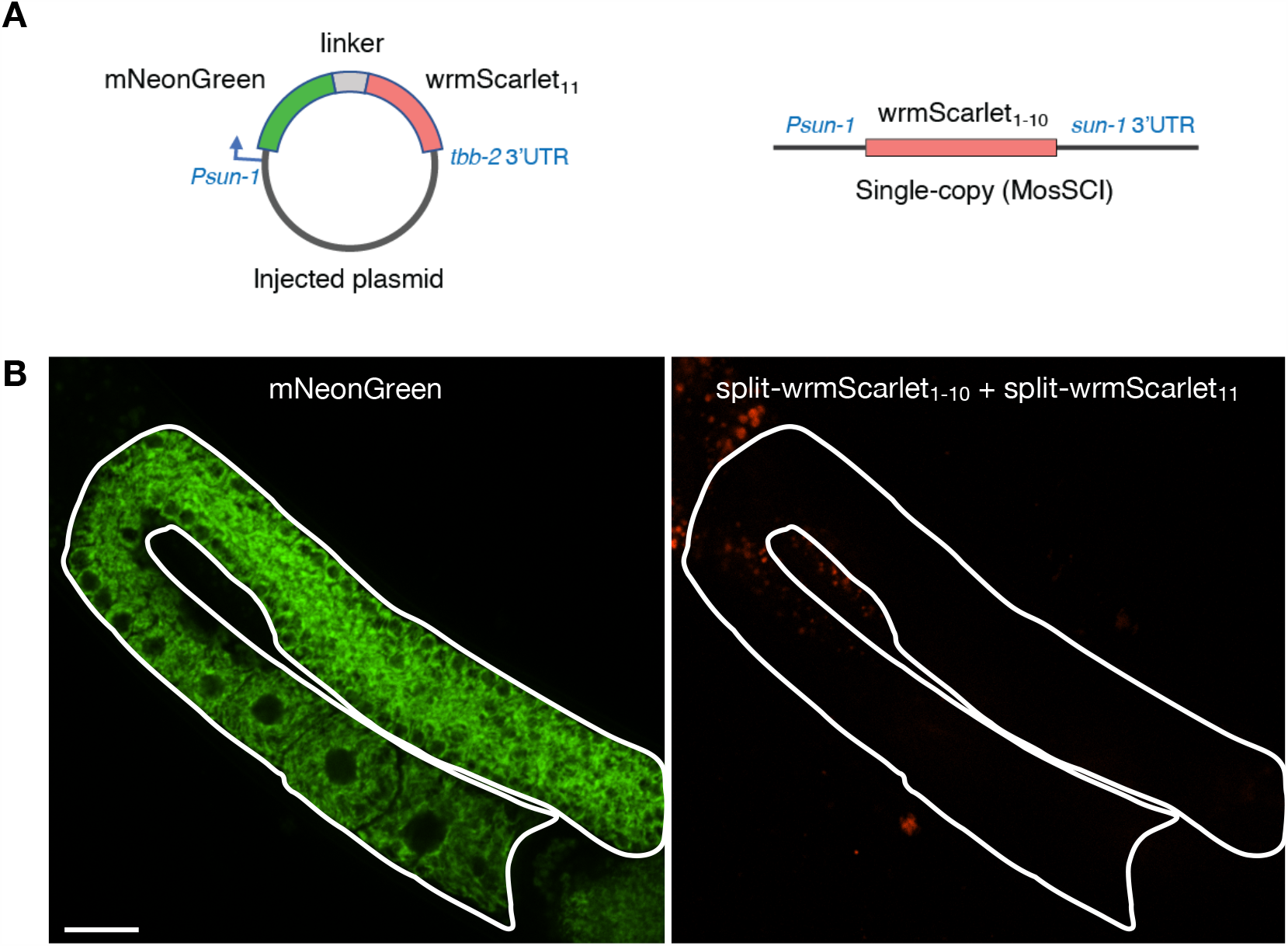
Tissue-specific split-wrmScarlet fluorescence in the germline is undetectable when split-wrmScarlet_1-10_ is integrated using a single-copy transgene via MosSCI. (A) Schematic of the plasmid encoding *Psun-1::mNeonGreen::linker::split-wrmScarlet*_*11*_*::tbb-2* 3’UTR (left), which was injected into the MosSCI strain PHX1797 carrying a single, integrated copy of *Psun-1::split-wrmScarlet*_*1-10*_*::sun-1* 3’UTR (right). (B) Images of animal expressing mNeonGreen::linker::split-wrmScarlet_11_ and split-wrmScarlet_1-10_ in the germline. Despite detecting mNeonGreen fluorescence, split-wrmScarlet was undetectable in this MosSCI strain, potentially due to compromised expression, folding or maturation of split-wrmScarlet_1-10_. Scale bar, 20 µm.

**Figure S8.**
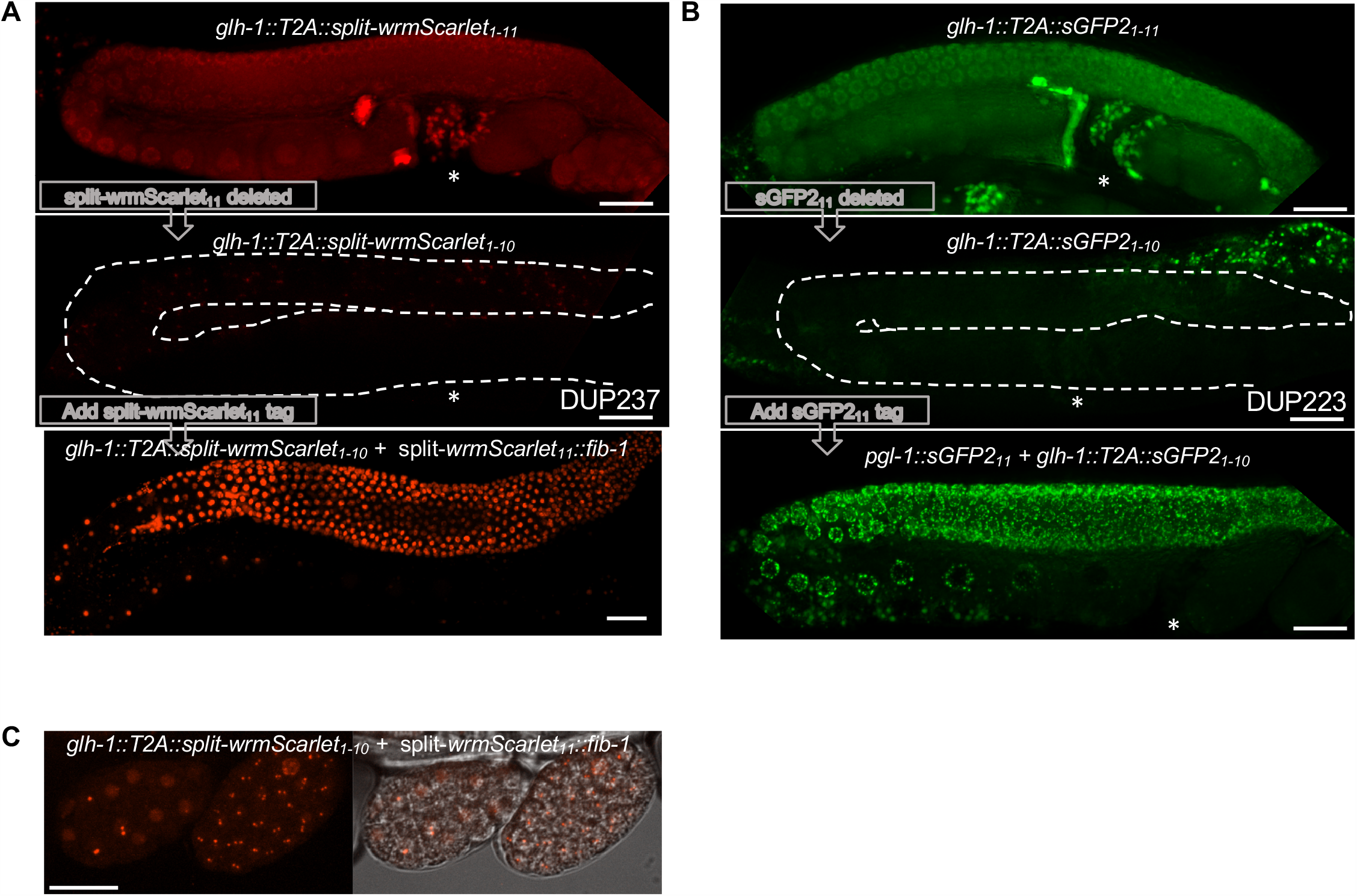
Generation and validation of germline-specific split-wrmScarlet_1-10_ and sGFP2_1-10_ strains. In order to generate germline-specific split-fluorescent strains, we first tagged the C-terminus of *glh-1* with T2A::split-wrmScarlet_1-11_ (A) or T2A::sGFP2_1-11_ (B). The T2A separates GLH-1 post-translationally to disperse the fluorophore throughout germ-cell nuclei, syncytium, sperm (*) and early embryos (Upper panels). We then deleted split-wrmScarlet_11_ or sGFP2_11_ to generate the corresponding split-FP_1-10_ strains DUP237 and DUP223 respectively. As expected, these strains were non-fluorescent (Middle panels). Tagging FIB-1 with split-wrmScarlet_11_, or PGL-1 with sGFP2_11_ confirmed that germline-specific labeling with split-wrmScarlet and split-sGFP2 can be successfully achieved using these strains (Lower panels). (C) Split-wrmScarlet_11_::FIB-1 is detectable in early embryos. Scale bars, 20 µm.

**Figure S9.**
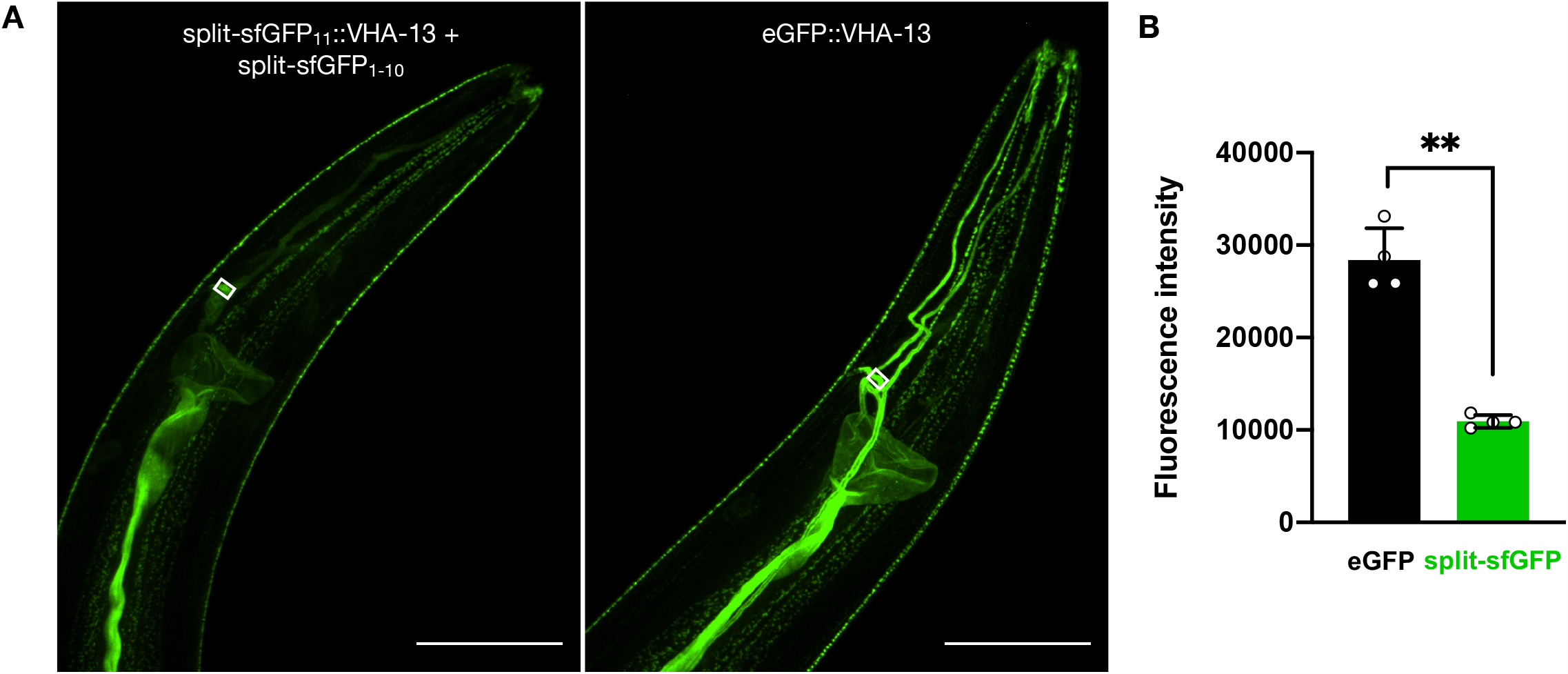
Somatic sfGFP_11_ compared to full-length eGFP at the endogenous *vha-13* locus. (A) Representative images of animals expressing sfGFP_1-10_ in somatic tissues with endogenous VHA-13 tagged with sfGFP_11_ (left panel), or endogenous VHA-13 tagged with eGFP in a wild-type background (right panel). Maximum intensity projections of 3D stacks shown. Scale bars, 50 µm. (B) Emission intensities from somatic sfGFP::VHA-13 and eGFP::VHA-13. Quantification was performed in the cell body, as quantifications in the excretory canal had higher variance. Mean ± s.d. Circles are individuals (n=4 for each condition). \*\**P* < 0.005.

**Figure S10.**
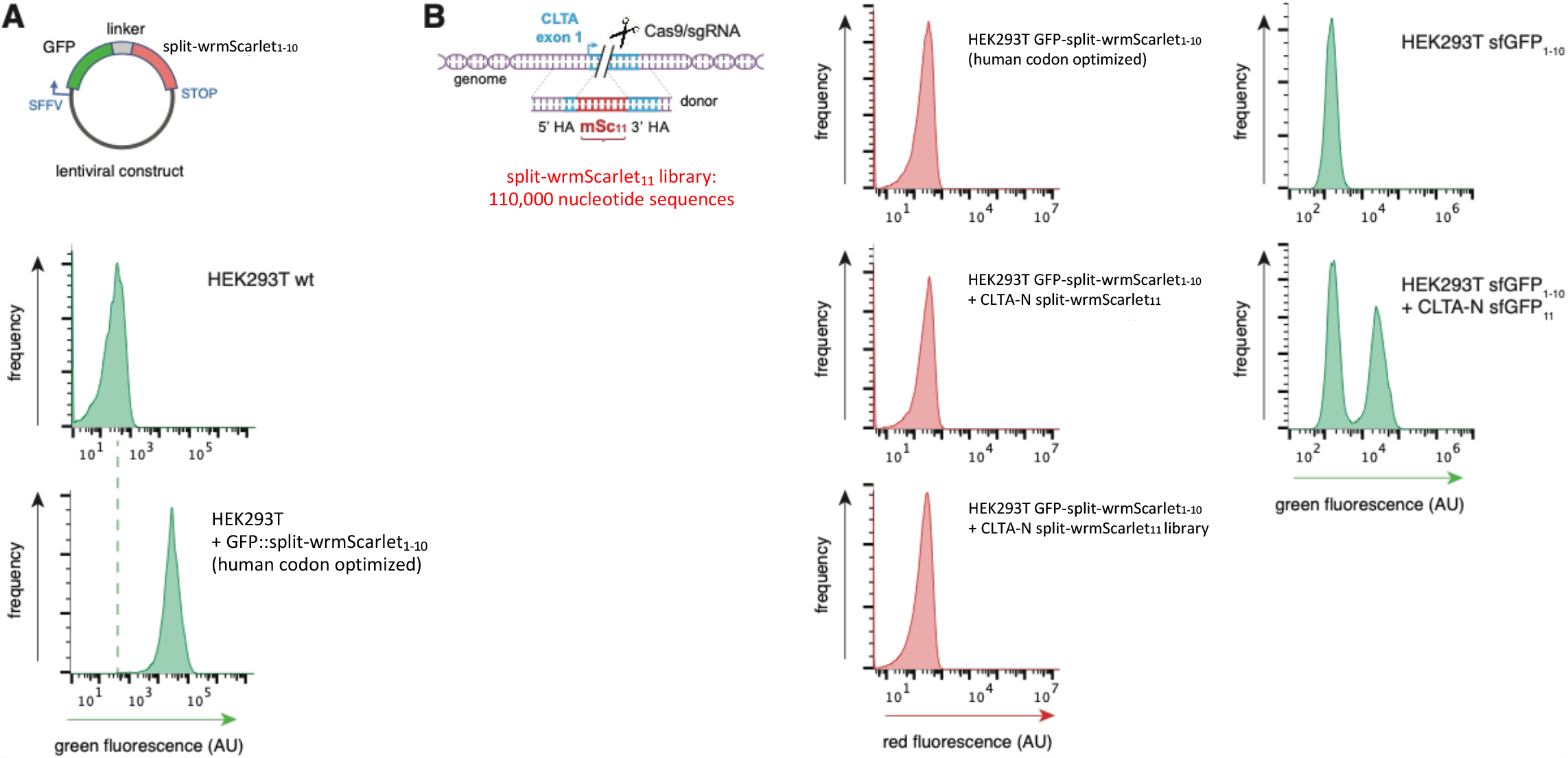
Screen for split-wrmScarlet fluorescence in mammalian cells. (A) FACS histograms of human codon-optimized split-wrmScarlet_1-10_ expressed as a C-terminal GFP fusion. GFP expression verifies successful expression of the fusion protein in HEK293T cells by lentiviral transduction. (B) Schematic of the CRISPR-based knock-in design for screening single and double mutants of split-wrmScarlet_11_. Left panel shows that neither our original split-wrmScarlet_11_ sequence nor its mutant library enabled detectable complementation as detected by FACS. Right panel shows that the control experiment using the sfGFP_1-10_/sfGFP_11_ system displays high levels of knock-in and complementation in HEK293T cells.

**Figure S11.**
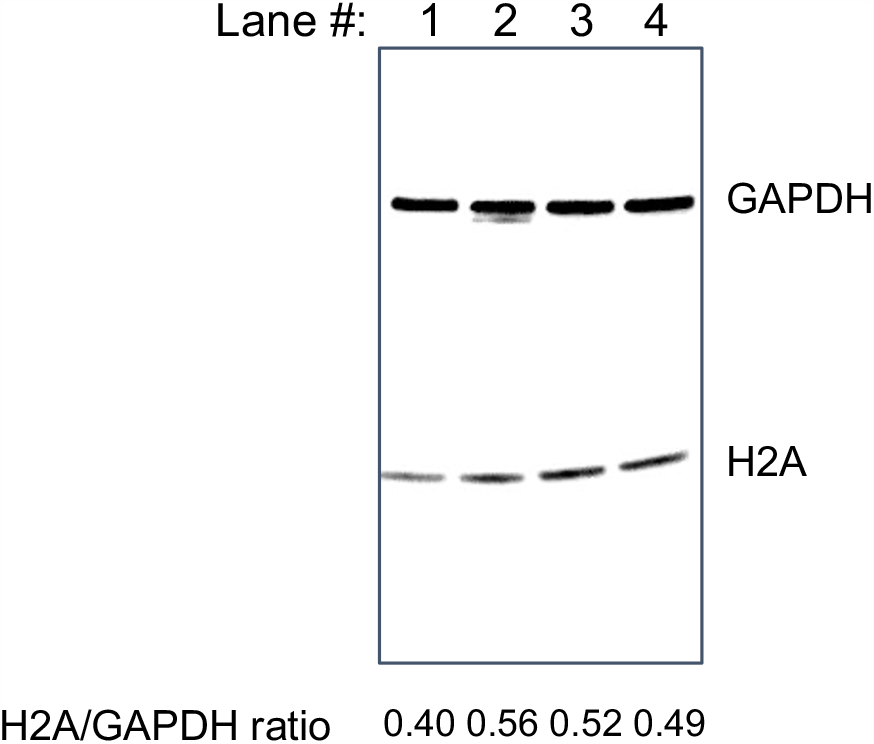
Split-wrmScarlet_11_ C-terminal amino acids did not affect H2A abundance. Western-blot of histone H2A (*his-3*) with split-wrmScarlet_11_+/- MDELYK. Western blot targeting HIS-3 in wild-type animals (lane 1), somatic split-wrmScarlet_1-10_ expressing animals (lane 2), somatic split-wrmScarlet_1-10_ strain with HIS-3::split-wrmScarlet_11_ ending with two glycines (lane 3), or somatic split-wrmScarlet_1-10_ + HIS-3::split-wrmScarlet_11(MDELYK)_. GAPDH was used as a loading control, and the HIS-3/GAPDH ratios are displayed under each lane.

**Figure S12.**
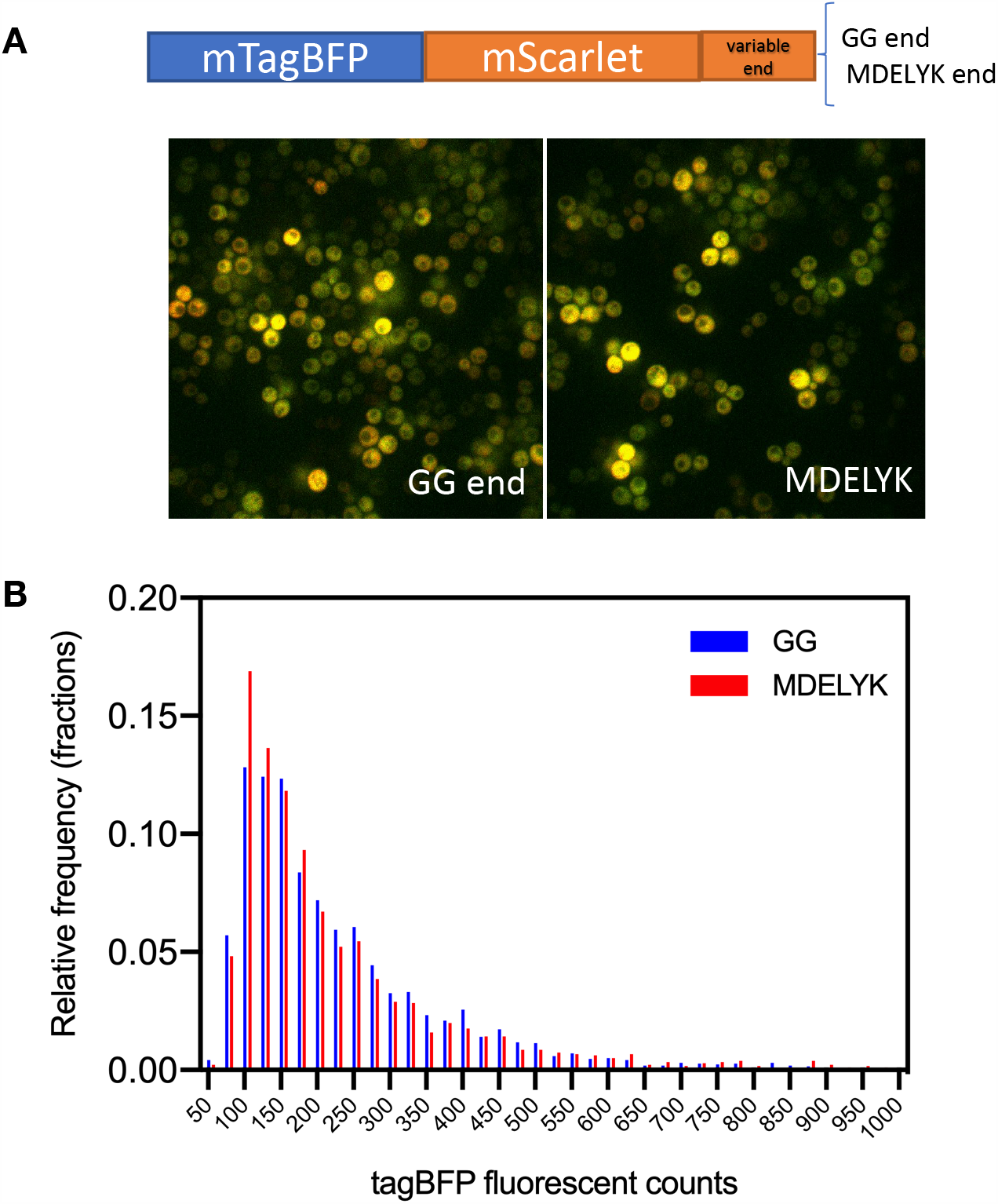
mScarlet ending with GG or MDELYK yields similar protein abundance in yeast. (A) Representative images of yeast expressing mTagBFP::mScarlet fusion truncated to end with gly-gly (GG end) or MDELYK from a p416-GPD promoter plasmid. mTagBFP fluorescence is pseudocolored in green, and mScarlet in red. (B) Histogram of mTagBFP fluorescence from 2556 yeasts expressing the truncation after GG and 1777 yeasts expressing the MDELYK end.

## SUPPLEMENTARY TEXT

### Supplementary Materials and Methods

#### Mammalian cell culture

HEK293T cells (ATCC # CRL-3216) were cultured in high-glucose DMEM supplemented with 10% FBS, 1 mM glutamine and 100 µg/mL penicillin/streptomycin (Gibco). An split-wrmScarlet_1-10_ cDNA codon-optimized for mammalian expression was fused to the C-terminus of eGFP and cloned into a pCDH lentiviral expression vector (SFFV GFP-mScarlet_1-10_). Lentivirus was prepared using standard protocols (Kamiyama *et al*. 2016) and used to infect HEK293T cells. A polyclonal population of GFP-split-wrmScarlet_1-10_ positive cells was isolated by FACS (using GFP fluorescence) and served as parental cell line for further experiments. For CLTA-N CRISPR engineering, *S. pyogenes* Cas9/sgRNA ribonucleoprotein complexes were prepared as in (Leonetti *et al*. 2016), mixed with HDR donor templates and electroporated into of GFP-split-wrmScarlet_1-10_ cells by nucleofection.

#### CLTA-N split-wrmScarlet11 donor library

A cDNA pool of degenerate split-wrmScarlet_11_ sequences was generated by oligonucleotide synthesis (GenScript) and homology arms for HDR-mediated insertion at CLTA N-terminus were appended by PCR (Supplementary Material - Table-S1 for full sequences). Library diversity was verified by Illumina MiSeq deep-sequencing.

## SUPPLEMENTARY RESULTS

### Split-wrmScarlet screening in mammalian cells

We tested the applicability of the split-wrmScarlet_1-10_ system for mammalian cell engineering but were surprisingly unsuccessful at detecting fluorescence. We designed a human codon-optimized split-wrmScarlet_1-10_ cDNA and expressed it as a C-terminal GFP fusion in HEK293T cells by lentiviral transduction. Expression of GFP verified the successful expression of the fusion protein (Figure S10A). However, subsequent expression of split-wrmScarlet_11_ fragments did not give rise to detectable red fluorescence despite numerous attempts. We reasoned that the split-wrmScarlet_11_ amino-acid sequence might be sub-optimal for complementation in human cells and synthesized a library of degenerate split-wrmScarlet_11_ sequences covering any possible single and double amino-acid mutants. Using an established assay for CRISPR-based knock-in of sequences at the CLTA N-terminus (a highly expressed gene in HEK293T cells (Leonetti *et al*. 2016), neither our original split-wrmScarlet_11_ sequence nor its mutant library enabled detectable complementation (Figure S10B, left panels). By contrast, a control experiment using the sfGFP_1-10_/sfGFP_11_ system showed a high level of knock-in and complementation in HEK293T (Figure S10B, right panels). It is possible that split-wrmScarlet_1-10_ is expressed in a non-functional form in human cells, or that its binding to split-wrmScarlet_11_ is occluded by competing interactions (with cellular chaperones, for example). In addition, we did not attempt complementation in primary non-transformed cell lines, like WI-38 cells, whose different proteostasis network and chaperones could aid split-wrmScarlet folding. At this point, more experiments would be required to establish the portability of split-wrmScarlet to mammalian systems.

### Experiments to investigate whether split-wrmScarlet_11_ functions as a degron in *C. elegans*

After finalizing our experiments, a paper that shows that C-terminal gly-gly sequences might function as degrons in mammalian cells was brought to our attention (Koren, *et al*. 2018). Since we were unable to obtain non-sterile positive clones of TOMM-20::split-wrmScarlet_11_ that did not have a mutation on the last glycine, and we were also unable to obtain EAT-6 homozygotes, we were concerned that our split-wrmScarlet_11_ might be recognized as a degron. To investigate this, we first labeled HIS-3, EAT-6, and TOMM-20 with the 24 a.a. split-wrmScarlet_11(MDELYK)_, which adds the sequence MDELYK to the C-terminus of split-wrmScarlet_11_. These worms were fertile, and at least as bright as those labeled with split-wrmScarlet_11_ (Figure S6). However, increased fluorescence could be due to increased molecular brightness rather than increased abundance. To address this, we compared the abundance of nuclear HIS-3, HIS-3::split-wrmScarlet_11_, and HIS-3::split-wrmScarlet_11(MDELYK)_ by western blot, and were unable to detect a significant change in abundance (Figure S11). We also could not detect differences in abundance in *S. cerevisiae*, using a p416-GPD plasmid expressing a mTagBFP-mScarlet fusion or the same fusion truncated so that it ends with gly-gly (Figure S12). However, because HIS-3 is a nuclear protein, and expression in yeast was done from an overexpressing plasmid, we cannot exclude that a protein ending with two glycines might be recognized as a degron in other cellular compartments, or at different expression levels, nor that there is no DesCEND degron pathway in yeast and worms. For these reasons, we recommend using the 24 a.a. split-wrmScarlet_11(MDELYK)_ when labeling proteins at their C-termini.

### Supplementary Materials – Supplementary Tables

**Table S1.**
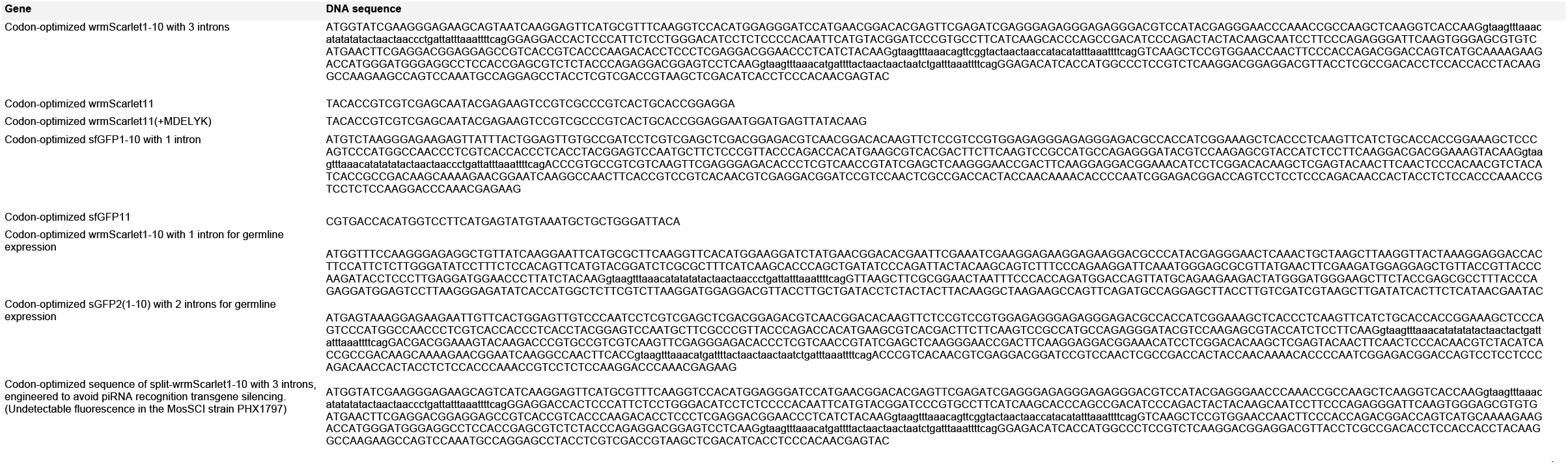
DNA Sequences of wrmScarletl −10, wrmScarletl 1, sfGFPI −10 and sfGFP11

**Table S2.**
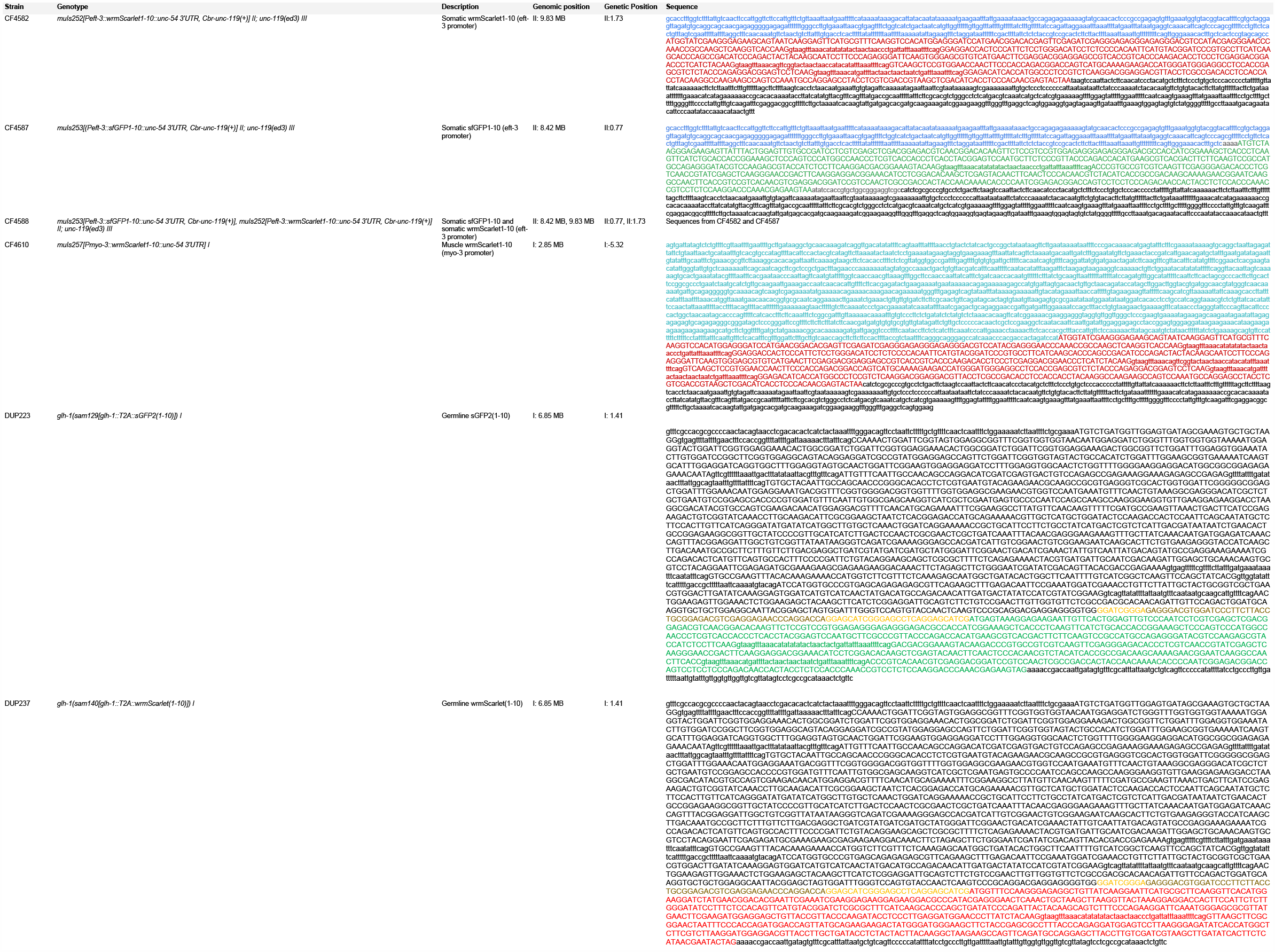
*C. elegans* l nes express ng s ngle-copy of wrm5carle 1-10 and/or sf FP1-10 5 ra n eno ype

**Table S3.**
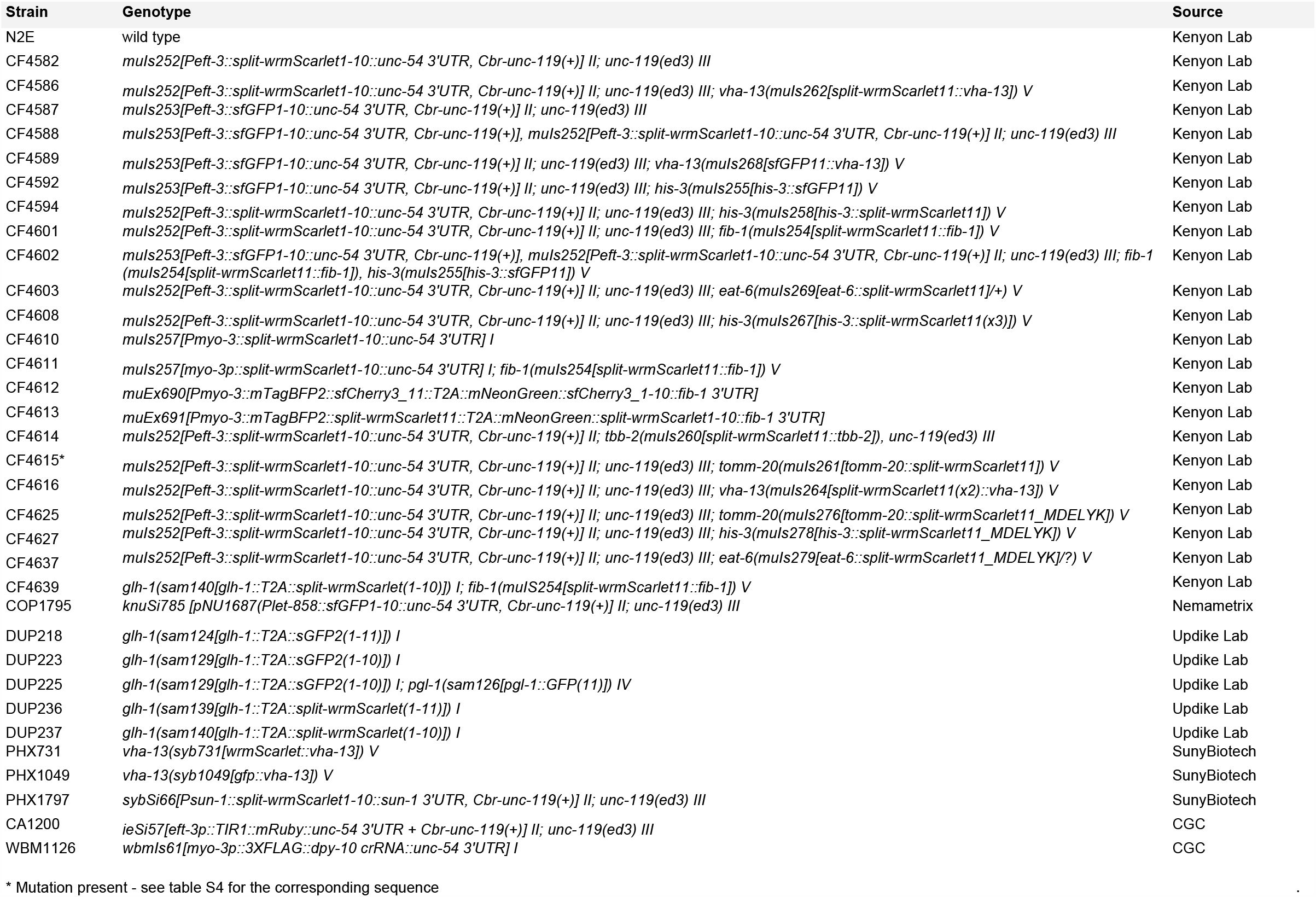
*C. elegans* strains, genotypes and sources

**Table S4.**
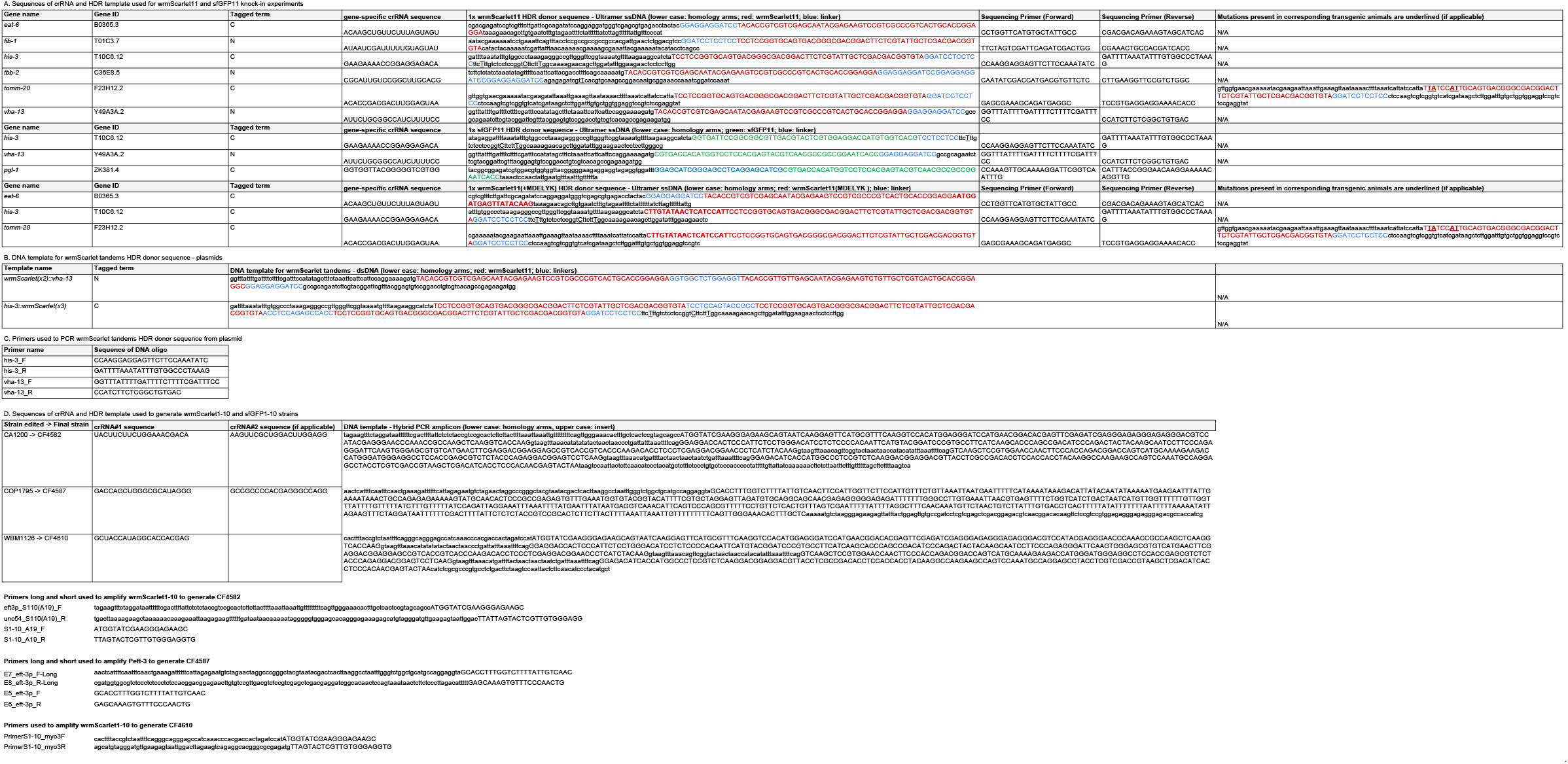
crRNAs, HDR templates and oligonucleotide sequences

**Table S5.**
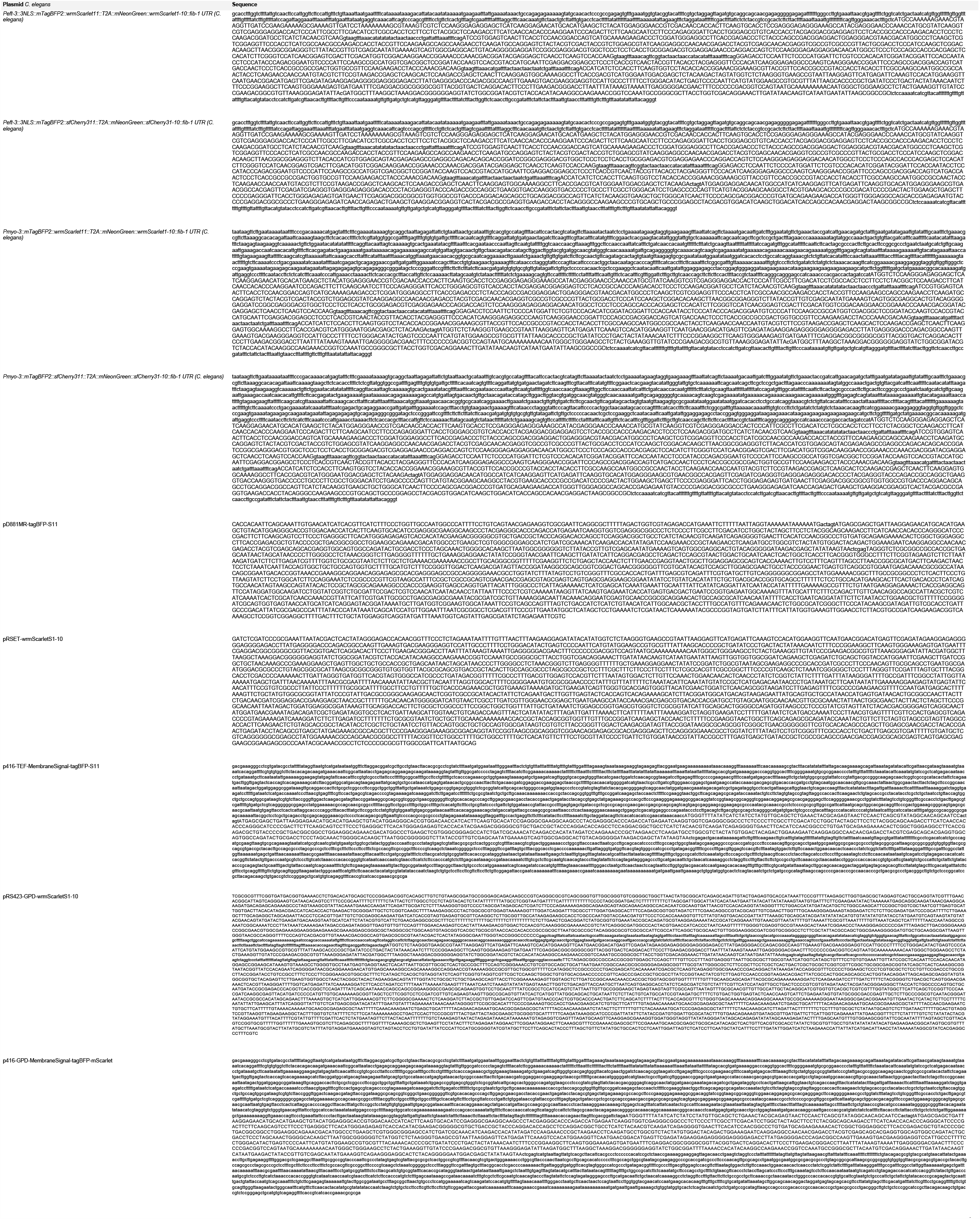
Plasmid sequences

**Table S6.**
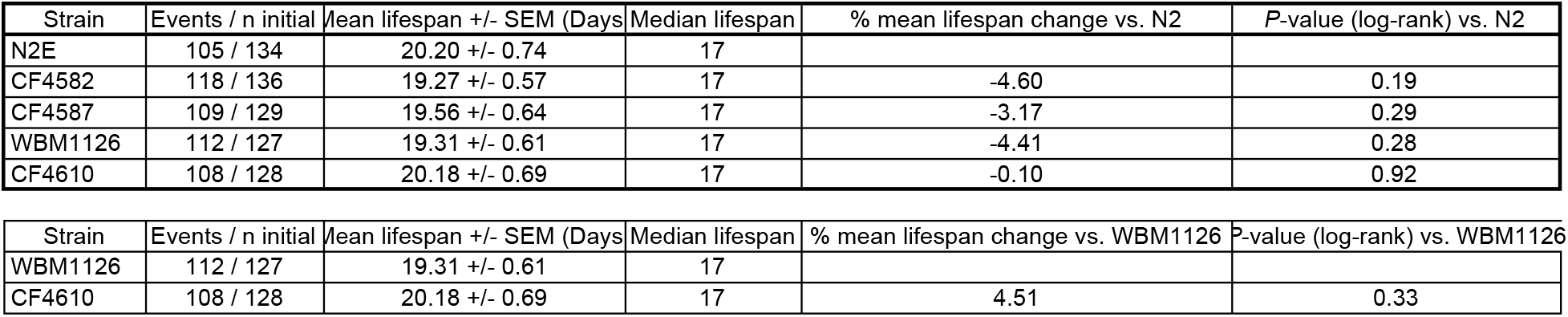
Adult lifespans data and statistics

## Step by step guide to tag endogenous genes with split-wrmScarlet and/or split-sfGFP in *C. elegans* V.1. doi.org/10.17504/protocols.io.bamkic4w

### MATERIALS

Microinjection practices and equipment, see WormBook chapter: www.wormbook.org/chapters/www_transformationmicroinjection/transformationmicroinjection.html

Evans, T. C., ed. Transformation and microinjection (April 6, 2006), WormBook, ed. The *C. elegans* Research Community, WormBook. doi.org/10.1895/wormbook.1.108.1, http://www.wormbook.org.

### BEFORE STARTING

If you are not already, become familiar with CRISPR/Cas genome editing, and general guidelines (Paix, *et al*. 2015; Kohler & Dernburg 2016; Farboud *et al*. 2019).

1. Select a *C. elegans* strain expressing split-wrmScarlet_1-10_ and/or sfGFP_1-10_ in your tissue of interest: Somatic split-sfGFP_1-10_: CF4587 *muIs253*[(*Peft-3::sfGFP*_*1-10*_*::unc-54 3’UTR, Cbr-unc-119*(*+*)] *II; unc-119*(*ed3*) *III* Somatic split-wrmScarlet_1-10_: CF4582 *muIs252*[*Peft-3::split-wrmScarlet*_*1-10*_*::unc-54 3’UTR, Cbr-unc-119*(*+*)] *II; unc-119*(*ed3*) *III* Dual Somatic split-sfGFP_1-10_ and split-wrmScarlet_1-10_: CF4588 *muIs253*[*Peft-3::sfGFP*_*1-10*_*::unc-54 3’UTR, Cbr-unc-119*(*+*)], *muIs252*[*Peft-3::split-wrmScarlet*_*1-10*_*::unc-54 3’UTR, Cbr-unc-119*(*+*)] *II; unc-119*(*ed3*) *III* Muscle-specific split-wrmScarlet_1-10_: CF4610 *muIs257*[*Pmyo-3::split-wrmScarlet*_*1-10*_*::unc-54 3’UTR*] *I* Germline-specific split-wrmScarlet_1-10_: DUP237 *glh-1*(*sam140*[*glh-1::T2A::split-wrmScarlet*_*1-10*_]) *I* Germline-specific sGFP2_1-10_: DUP223 *glh-1*(*sam129*[*glh-1::T2A::sGFP2*_*1-10*_]) *I*
2. Select a guide sequence and order a crRNA and tracrRNA. Download the DNA sequence of your gene and transcript of interest from Wormbase. Identify the desired insertion or knock-in site in the genomic DNA.
3. Using ∼50 nucleotides flanking Identify DNA sequence(s) followed by a PAM site (5’ NGG 3’) in your target gene using a CRISPR/Cas9 target online predictor, such as CCTop (Stemmer *et al*. 2015; Labuhn, M. *et al*. 2018). If using CCTop, use the default parameters and adjust the following two criteria: PAM type: NGG (*Streptococcus pyogenes*) Species: *C. elegans* Insert the DNA sequence in the query section, then submit. Select the crRNA target site with a high score, the closest to your editing site, with no off-targets.
4. Order the crRNA corresponding to your selected guide sequence from IDT. Note: Do not include the PAM in the query. Typically, we order 10 nmol of each Alt-R® CRISPR-Cas9 crRNA that we resuspend in 14.5 µL TE.
5. Order the universal 67 mer tracrRNA from IDT under the section “CRISPR-Cas9 tracrRNA” (www.idtdna.com/pages/products/crispr-genome-editing/alt-r-crispr-cas9-system). Typically, we order 20 nmol.
6. Design and order single-stranded donor oligonucleotides (ssODN) of 200-mers. Design the sequence of a single-stranded donor oligonucleotides (ssODN) with the Fluorescent Protein_11_ of choice, (*i*.*e*. split-wrmScarlet_11_ or sfGFP_11_), a linker and homology arms. Sequences: Split-wrmScarlet_11_ for N-terminus labeling: 5’ TACACCGTCGTCGAGCAATACGAGAAGTCCGTCGCCCGTCACTGCACCGGAGGA 3’ Split-wrmScarlet_11_ for C-terminus labeling: 5’ TACACCGTCGTCGAGCAATACGAGAAGTCCGTCGCCCGTCACTGCACCGGAGGAATGGATGAGTTATACAAG 3’ sfGFP_11_: 5’ CGTGACCACATGGTCCTTCATGAGTATGTAAATGCTGCTGGGATTACA 3’ Linker: 5’ GGAGGAGGATCC 3’ The linker should be inserted at the 5’ end or 3’ end of the FP_11_ depending on the site chosen to insert it in your gene of interest, for example:
  - When tagging the N-terminal of your gene of interest, the final ssODN sequence should look like this: Left homology arm :: endogenous START codon (ATG) :: Fluorescent Protein_11_ :: Linker :: Right homology arm
  - When tagging the N-terminal of a gene of interest, the final ssODN sequence should look like this:
  Left homology arm :: Linker:: Fluorescent Protein_11_ :: endogenous STOP codon :: Right homology arm Ensure that the linker and the Fluorescent Protein_11_ sequences are in frame with the coding sequence of your gene of interest. Linker length and flexibility may need to be optimized depending on the protein structure and function. The final ssODN can be up to 200 mers, allowing each homology arms to be 67 nucleotides when tagging a gene with split-wrmScarlet_11_ and the suggested linker at the N-terminus or 58 for tagging at the C-terminus. Order the ssODN from IDT in the section “Ultramer DNA Oligos”. We typically order up to 4 nmol at 100 µM in IDTE buffer
7. Design and order oligos that will be used to perform the genotyping and sequencing of the insertion site.
  - 18-24 bases in length
  - Melting temperature (Tm) between 50 and 60°C
  - GC content ranging from 45 to 55%
  - G or C on the 3’ end
  - Design primers 200 nt upstream and downstream from the sequence of interest
8. Prepare and inject CRISPR/Cas9 ribonucleoprotein mix Assemble CRISPR/Cas9 ribonucleoproteins complex and ssODN into injection mix: <u>Protocol without co-CRISPR</u> 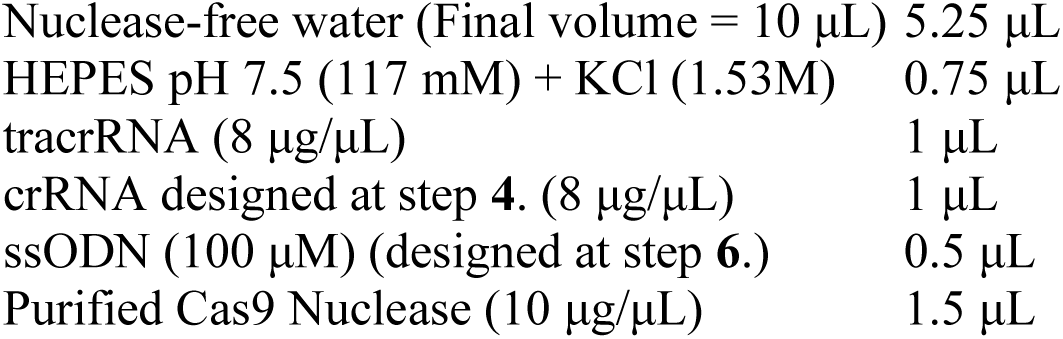 <u>Protocol with co-CRISPR</u>*<u>dpy-10</u>*(*<u>cn64</u>*) 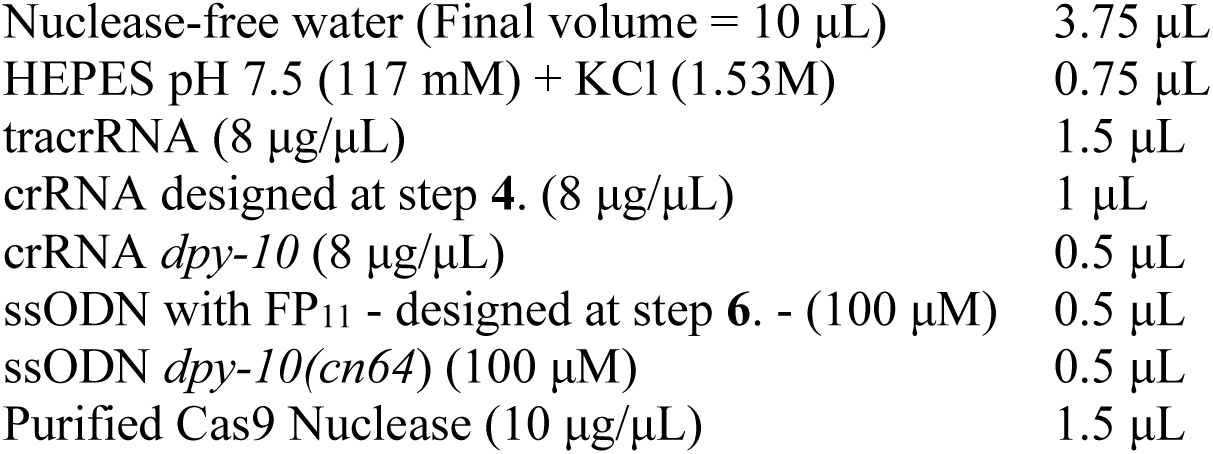 Pipet up and down a few times without introducing air into the mix. Incubate 37 °C 15 min Quick spin 13000 rpm Load 0.5 to 1 µL of the mix in injection needles and inject 10 to 20 day-1 adult worms successfully, ideally in both gonad arms. Single injected worms in a drop of M9 buffer on NGM plates seeded with OP50, and place them in a 25 °C incubator Overnight.
9. Screen for successful FP_11_ integrants Starting 3 days post injections, screen daily until identifying fluorescent progeny in the F1 and/or F2 of singled injected worms. Perform PCR genotyping and sequencing using regular worm protocols with the primers designed in step **7**.

## Literature cited

Bindels, D. S. et al. mScarlet: a bright monomeric red fluorescent protein for cellular imaging. Nature Publishing Group 14, 53–56 (2016).

Brenner, S. The genetics of Caenorhabditis elegans. Genetics 77, 71–94 (1974).

Cabantous, S., Terwilliger, T. C. & Waldo, G. S. Protein tagging and detection with engineered self-assembling fragments of green fluorescent protein. Nat Biotechnol 23, 102–107 (2004).

Dickinson, D. J. & Goldstein, B. CRISPR-Based Methods for Caenorhabditis elegans Genome Engineering. Genetics 202, 885–901 (2016).

Dokshin, G. A., Ghanta, K. S., Piscopo, K. M. & Mello, C. C. Robust Genome Editing with Short Single-Stranded and Long, Partially Single-Stranded DNA Donors in Caenorhabditis elegans. Genetics 210, 781–787 (2018).

El Mouridi, S. et al. Reliable CRISPR/Cas9 Genome Engineering in Caenorhabditis elegans Using a Single Efficient sgRNA and an Easily Recognizable Phenotype. G3 7, 1429–1437 (2017).

Evans, T. C., ed. Transformation and microinjection (April 6, 2006), WormBook, ed. The C. elegans Research Community, WormBook, doi: 10.1895/wormbook.1.108.1, http://www.wormbook.org.

Farboud, B., Severson, A. F. & Meyer, B. J. Strategies for Efficient Genome Editing Using CRISPR-Cas9. Genetics 211, 431–457 (2019).

Feng, S. et al. Improved split fluorescent proteins for endogenous protein labeling. Nature Communications 8, 370 (2017).

Feng, S. et al. Bright split red fluorescent proteins for the visualization of endogenous proteins and synapses. Communications Biology 2, 344–12 (2019).

Friedland, A. E. et al. Heritable genome editing in C. elegans via a CRISPR-Cas9 system. Nature Publishing Group 10, 741–743 (2013).

Frøkjær-Jensen, C., Davis, M. W., Ailion, M. & Jorgensen, E. M. Improved Mos1-mediated transgenesis in C. elegans. Nature Publishing Group 9, 117–118 (2012).

Frøkjær-Jensen, C. et al. Single-copy insertion of transgenes in Caenorhabditis elegans. Nat Genet 40, 1375–1383 (2008).

Goudeau, J. et al. Fatty acid desaturation links germ cell loss to longevity through NHR-80/HNF4 in C. elegans. PLoS Biol 9, e1000599 (2011).

He, S., Cuentas-Condori, A. & Miller, D. M. NATF (Native and Tissue-Specific Fluorescence): A Strategy for Bright, Tissue-Specific GFP Labeling of Native Proteins in Caenorhabditis elegans. Genetics 212, 387–395 (2019).

Hefel, A. & Smolikove, S. Tissue-Specific Split sfGFP System for Streamlined Expression of GFP Tagged Proteins in the Caenorhabditis elegans Germline. G3 9, 1933–1943 (2019).

Hu, C. D., Chinenov, Y. & Kerppola TK. Visualization of interactions among bZIP and Rel family proteins in living cells using bimolecular fluorescence complementation. Molecular Cell 9, 789–98 (2002).

Ramadani-Muja, J. et al. Visualization of Sirtuin 4 Distribution between Mitochondria and the Nucleus, Based on Bimolecular Fluorescence Self-Complementation. Cells 8, (2019).

Kamiyama, D. et al. Versatile protein tagging in cells with split fluorescent protein. Nature Communications 7, 11046–9 (2016).

Kelley, L. A., Mezulis, S., Yates, C. M., Wass, M. N. & Sternberg, M. J. E. The Phyre2 web portal for protein modeling, prediction and analysis. Nat Protoc 10, 845–858 (2015).

Kelly, W. G., Xu, S., Montgomery, M. K. & Fire, A. Distinct Requirements for Somatic and Germline Expression of a Generally Expressed Caenorhabditis elegans Gene. Genetics 146, 227–238 (1997).

Kimble, J., Hodgkin, J., Smith, T. & Smith, J. Suppression of an amber mutation by microinjection of suppressor tRNA in C. elegans. Nature 299, 456–458 (1982).

Köker, T., Fernandez, A. & Pinaud, F. Characterization of Split Fluorescent Protein Variants and Quantitative Analyses of Their Self-Assembly Process. Sci. Rep. 8, 5344 (2018).

Koren I, Timms RT, Kula T, Xu Q, Li MZ, Elledge SJ. The Eukaryotic Proteome Is Shaped by E3 Ubiquitin Ligases Targeting C-Terminal Degrons. Cell. 2018 Jun 14;173(7):1622-1635.e14. doi: 10.1016/j.cell.2018.04.028. Epub 2018 May 17. PMID: 29779948; PMCID: PMC6003881.

Liu, Z., Chen, O., Wall, J., Zheng, M., Zhou, Y., Wang, L., Vaseghi, H. R., Qian, L., & Liu, J. (2017). Systematic comparison of 2A peptides for cloning multi-genes in a polycistronic vector. Scientific reports, 7(1), 2193. https://doi.org/10.1038/s41598-017-02460-2.

Leonetti, M. D., Sekine, S., Kamiyama, D., Weissman, J. S. & Huang, B. A scalable strategy for high-throughput GFP tagging of endogenous human proteins. Proc. Natl. Acad. Sci. U.S.A. 113, E3501–8 (2016).

Marnik, E. A. et al. Germline Maintenance Through the Multifaceted Activities of GLH/Vasa in Caenorhabditis elegans P Granules. Genetics 213, 923–939 (2019).

Mello, C. C., Kramer, J. M., Stinchcomb, D. & Ambros, V. Efficient gene transfer in C. elegans: extrachromosomal maintenance and integration of transforming sequences. The EMBO Journal 10, 3959–3970 (1991).

Nance, J. & Frøkjær-Jensen, C. The Caenorhabditis elegans Transgenic Toolbox. Genetics 212,959–990 (2019).

Noma, K., Goncharov, A., Ellisman, M. H. & Jin, Y. Microtubule-dependent ribosome localization in C. elegans neurons. eLife 6, 218 (2017).

Paix, A. et al. Scalable and versatile genome editing using linear DNAs with microhomology to Cas9 Sites in Caenorhabditis elegans. Genetics 198, 1347–1356 (2014).

Paix, A. High Efficiency, Homology-Directed Genome Editing in Caenorhabditis elegans Using CRISPR-Cas9 Ribonucleoprotein Complexes. 1–26 (2015). doi:10.1534/genetics.115.179382/-/DC1.

Paix, A., Schmidt, H. & Seydoux, G. Cas9-assisted recombineering in C. elegans: genome editing using in vivo assembly of linear DNAs. Nucleic Acids Res 44, e128 (2016).

Pédelacq, J.-D., Cabantous, S., Tran, T., Terwilliger & T. C., Waldo, G. S. Engineering and characterization of a superfolder green fluorescent protein. Nature Biotechnology 24(1), 79–88 (2005).

Prior, H., Jawad, A. K., MacConnachie, L. & Beg, A. A. Highly Efficient, Rapid and Co-CRISPR-Independent Genome Editing in Caenorhabditis elegans. G3 7, 3693–3698 (2017).

Redemann, S. et al. Codon adaptation-based control of protein expression in C. elegans. Nature Publishing Group 8, 250–252 (2011).

Richardson, C. D. et al. CRISPR-Cas9 genome editing in human cells occurs via the Fanconi anemia pathway. Nat Genet 50, 1132–1139 (2018).

Silva-García, C. G. et al. Single-Copy Knock-In Loci for Defined Gene Expression in Caenorhabditis elegans. G3 9, 2195–2198 (2019).

Snapp, E. Design and use of fluorescent fusion proteins in cell biology. Current Protocols in Cell Biology Chapter 21, 21.4.1-21.4.13 (2005).

Vicencio, J., Martínez-Fernández, C., Serrat, X. & Cerón, J. Efficient Generation of Endogenous Fluorescent Reporters by Nested CRISPR in Caenorhabditis elegans. Genetics 211, 1143–1154 (2019).

Wu, W.-S. et al. pirScan: a webserver to predict piRNA targeting sites and to avoid transgene silencing in C. elegans. Nucleic Acids Res 46, W43–W48 (2018).

Zhang, L., Ward, J. D., Cheng, Z. & Dernburg, A. F. The auxin-inducible degradation (AID) system enables versatile conditional protein depletion in C. elegans. Development 142, 4374–4384 (2015).

Zhang, D. et al. The piRNA targeting rules and the resistance to piRNA silencing in endogenous genes. Science 359, 587–592 (2018).

## Supplementary Protocol – Literature cited

Kohler S, Dernburg A. (2016) C. elegans injection: Ribonucleoprotein delivery using the Alt-R CRISPR-Cas9 System. [Online] Coralville, Integrated DNA Technologies. [December, 2017.]

Labuhn, M. et al. Refined sgRNA efficacy prediction improves large- and small-scale CRISPR-Cas9 applications. Nucleic Acids Res 46, 1375–1385 (2018).

Paix et al. Direct delivery CRISPR-HDR editing protocol for C. elegans. http://dx.doi.org/10.17504/protocols.io.dri54d.

Paix, A., Folkmann, A., Rasoloson, D. & Seydoux, G. High Efficiency, Homology-Directed Genome Editing in Caenorhabditis elegans Using CRISPR-Cas9 Ribonucleoprotein Complexes. Genetics 201, 47–54 (2015).

Stemmer, M. et al. CCTop: An Intuitive, Flexible and Reliable CRISPR/Cas9 Target Prediction Tool. PLoS ONE 10, e0124633 (2015).

